# Axon Collateral Pattern of a Sparse Locus Coeruleus Norepinephrine Neuron in Mouse Cerebral Cortex

**DOI:** 10.1101/2025.02.04.636453

**Authors:** Yu-Ming Wu, Wei-Chen Hung, Yun Chang, Ming-Yuan Min, Hsiu-Wen Yang

## Abstract

The locus coeruleus (LC) contains predominantly norepinephrine (NE) neurons that project widely throughout the brain. The LC plays a critical role in controlling behavior, particularly arousal. Historically, it was thought that the LC-NE system performed its behavioral control function by uniformly releasing NE throughout most brain regions. However, recent evidence suggests that the LC’s cortical projections are organized into modules, which allows for the coordination of diverse, and sometimes opposing, functions such as fear memory formation and extinction. Nevertheless, many details remain unclear and require data from the axon collaterals of sparse neurons. We modified a viral tracing protocol using a dual-recombinase system to trace the axonal collaterals of sparse LC neurons projecting to the cingulate cortex (CgC). Our results show that even a small number of LC neurons have broad cortical projections, though the pattern is not uniform. Centered-log ratio transformation of NE fiber distribution across the cortex and hippocampus reveals a few preferential target areas (PTAs) of the labeled LC-NE neurons’ axonal projections. The summed NE fiber length in these defined PTs is enriched relative to the geometric mean of all other cortical and hippocampal regions where NE fibers were detected. Notably, the defined PTAs—including the rostral splenial cortex, dorsal hippocampus, somatosensory cortex, and CgC (the retrograde viral labeling injection site)—are functionally related to navigation. These results demonstrate that LC-NE neurons are organized into distinct projection modules, each comprising a small number of neurons with functionally correlated major cortical targets.

## Introduction

The locus coeruleus (LC) is composed primarily of norepinephrine (NE) neurons. Due to its widespread projections throughout the central nervous system, the LC serves as the primary source of NE in the brain and spinal cord (Schwarz et al., 2015; Aston-Jones & Waterhouse, 2016) and plays a pivotal role in behavioral control (Aston-Jones & Cohen, 2005; Bouret & Sara, 2005; Sales et al., 2019; Poe et al., 2020). LC-NE activity strongly modulates arousal (McCarley & Hobson, 1975; Aston-Jones & Bloom, 1981; Takahashi et al., 2010; Carter et al., 2010; Joshi et al., 2016; Lovett-Barron et al., 2017; Breton-Provencher & Sur, 2019; Hayat et al., 2020). Low firing rates are associated with sleep, moderate rates with wakefulness, and abnormally high rates with hyperactivity, attention deficits, and anxiety (Aston-Jones & Cohen, 2005). LC-NE neurons were traditionally thought to function via a uniform NE fiber projection to the cortex (Morrison et al., 1978; Biegon & Rainbow, 1983; Audet et al., 1988), enabling widespread NE release and simultaneous influence over entire neural networks in the cerebral cortex (Poe et al., 2020). However, recent findings regarding the pattern of axonal projections from the LC to the cortex and its role in behavioral regulation have challenged this view (Hansen & Manahan-Vaughan, 2015; Uematsu et al., 2017; Hirschberg et al., 2017; Takeuchi et al., 2016; McCall et al., 2017; Poe et al., 2020).

Quantitative analyses demonstrate that NE innervation is regionally specific, not uniform. For example, there are denser NE fibers in the prefrontal cortex than in the motor, somatosensory, and piriform cortices (Agster et al., 2013). Furthermore, trans-synaptic pseudorabies virus tracing suggests that LC-NE neurons exhibit differential subcircuit specificity (Schwarz et al., 2015). Retrograde virus tracing of axonal branches to the medial prefrontal cortex (mPFC) and primary motor cortex (M1) indicates distinct axon collateralization patterns (Uematsu et al., 2017; Plummer et al., 2020).

These findings are consistent with functional studies showing that LC-NE subcircuits orchestrate opposing learning states—facilitatory and extinctive fear learning—via distinct amygdala- and mPFC-projecting modules (Uematsu et al., 2017). Similarly, a spinal-projecting module promotes analgesia in acute pain, whereas a prefrontal-projecting module generates aversion in chronic pain (Hirschberg et al., 2017). In summary, the LC-to-cortex projection is highly structured into functional modules. Understanding this modular arrangement provides insight into how the LC-NE system regulates multiple functions to ensure adaptive responses and behavioral flexibility.

However, the details of the LC-NE organization remain unclear. Specifically, it is unclear if the LC-to-cortex projection comprises numerous LC-NE neurons, each with a specific cortical target, or if individual neurons have divergent projections to functionally correlated cortical regions that orchestrate different brain functions. To address this issue, data on the entire set of axonal collaterals from a sparse number of neurons, or even a single neuron, is necessary. This study aims to achieve this goal using the modified Tracing Axon Collaterals (TrAC) method (Plummer et al., 2020). The cingulate cortex (CgC) of the medial prefrontal cortex (mPFC) was targeted for retrograde virus tracing due to its critical role in emotional and cognitive functions (O’Doherty et al., 2002; Ito et al., 2003; Sanfey et al., 2003; Walton et al., 2007; Rolls, 2023), as well as its high noradrenaline (NE) fiber density compared to other cortical areas (Agster et al., 2013; Schwarz et al., 2015; Plummer et al., 2020). Furthermore, CgC function is largely modulated by the LC-NE system in complex tests, such as go/no-go (Bouret & Sara, 2005) and two-alternative forced-choice (2-AFC; Aston-Jones & Cohen, 2005). We found that even a few LC-NE neurons project broadly across the cortex; however, the pattern is not uniform. Using the centered log-ratio (CLR) transformation, we identified several preferential target areas (PTAs) with enriched NE fiber length relative to the geometric mean of all other areas. Importantly, the defined PTAs are functionally related cortical areas. Additionally, there were cortical areas with NE fibers that were defined as non-PTAs because the length of NE fibers within these areas was proportionate or depleted relative to the geometric mean. These non-PTAs are less functionally specific. We propose that the LC-NE system regulates arousal by projecting to these non-PTAs.

## Materials and Methods

### Experimental design for tracing axonal collaterals of sparse LC-NE neurons

The TrAC method described by Plummer et al. (2020) effectively traces complete axonal collaterals of LC-NE neurons. However, labeling too many neurons (up to hundreds) in a single experiment may obscure parts of their axonal collaterals due to overlap, leading to resolution loss. The TrAC method primarily uses a mouse strain carrying a dual recombinase-responsive fluorescent tracer (FT) allele crossed with a dopamine beta-hydroxylase (DBH, a synthesizing enzyme of NE and thus a producer of NE neurons) Flp mouse strain. This ensures that NE neurons in the offspring express both Flp recombinase and the FT allele. A second recombinase, Cre, is introduced into a subset of LC-NE neurons projecting to the cortical area of interest by retrograde canine viral infusion. This process activates FT expression, allowing visualization of the entire axonal collaterals of all labeled LC-NE neurons.

The number of labeled neurons depends on the amount of retrograde virus infused into the cortex and the infection efficiency of the viral vector. Since viral transfection rates are typically less than 100%, we refined this approach using dual AAV infusions while maintaining genetic specificity with dual recombinase-dependent FT expression. Specifically, we used tyrosine hydroxylase (TH, a marker for NE neurons due to its role as the rate-limiting enzyme for NE synthesis) Cre mice (strain #: 025614, Jackson Laboratory) in which the TH promoter drives Cre expression. We labeled a subset of LC-NE neurons projecting to the right CgC by infusing rAAV for Cre-dependent Flp expression into the cortical region (Figure 1A). Instead of tracking all retrogradely labeled LC-NE neurons as in the original TrAC method, we tracked a smaller subset by infusing AAV for Flp-dependent GCaMP6s (as FT) expression into the bilateral LC (Figure 1B). This dual AAV injection strategy successfully labeled sparse LC-NE neurons, allowing detailed tracing of their axonal collaterals.

**Figure 1.**
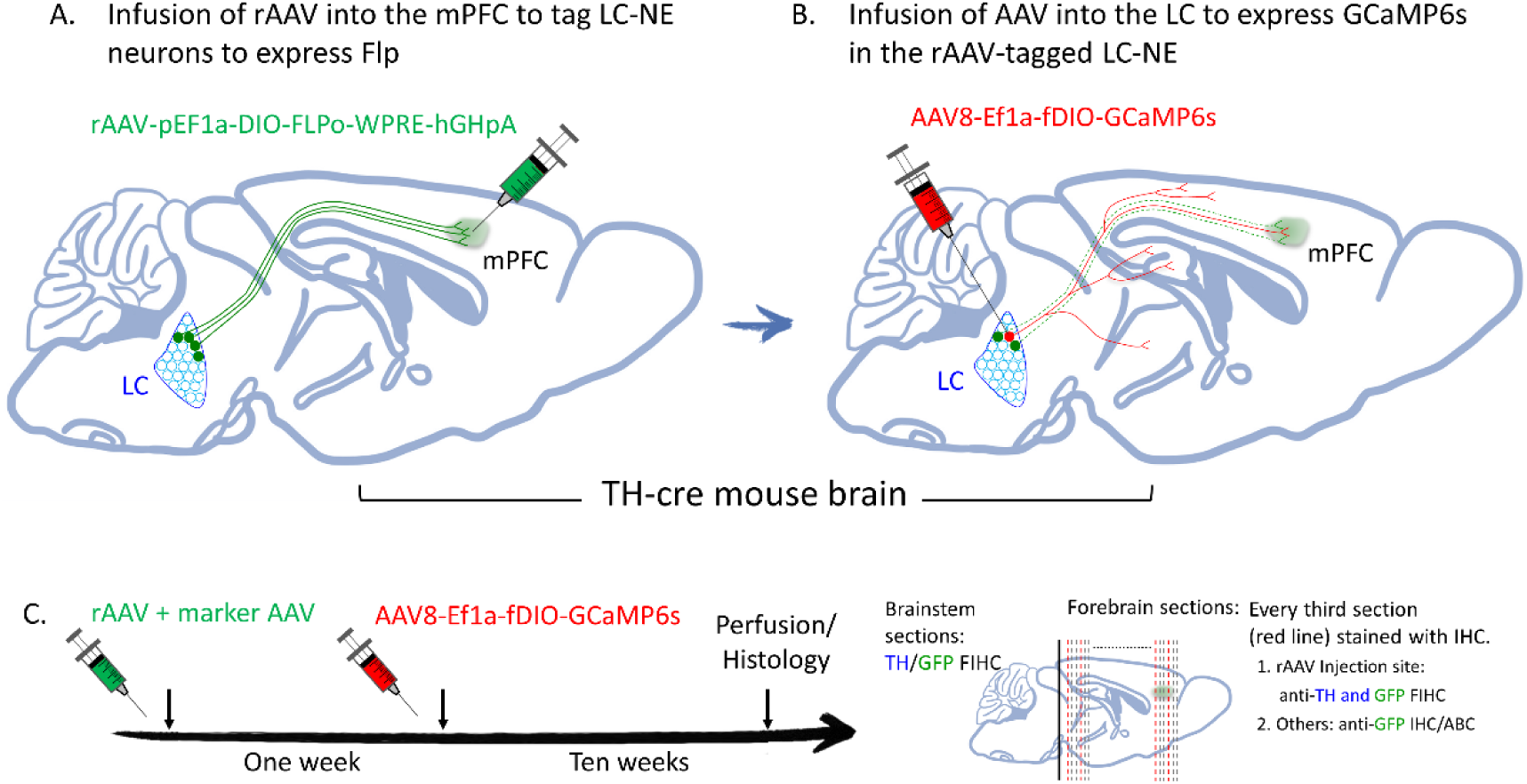
Modified TrAC to reveal the entire axon collateral of a sparse subset of LC-NE neurons. ***A***. Schematic illustration of retrograde AAV-mediated transduction of a subset of LC-NE neurons projecting to the CgC to express Flp. ***B***. A schematic illustrating the infusion of AAV8 into the LC to express GCaMP6s in a fraction of the rAAV-tagged LC-NE neurons that project to the CgC. ***C***. Timeline of rAAV and AAV8 infusions (left) and a schematic illustration of the blocking (thick line) of the brainstem and forebrain for sectioning after perfusion, as well as the grouping of the forebrain sections for IHC and Nissl staining (right).

### Animals and stereotactic surgery for viral vector injection

Mice of both sexes were used in all experiments. They were housed and maintained in a temperature-controlled vivarium under a 12-hour light/dark cycle, with food and water available *ad libitum*. All animal experiments were approved by the Institutional Animal Care and Use Committees, and every effort was made to minimize the number of animals used and their suffering. For stereotaxic surgery, 4-to 6-week-old mice were deeply anesthetized, as indicated by the absence of the hind paw withdrawal reflex, with an intraperitoneal injection of a mixture of zoletil (25 mg/kg) and xylazine hydrochloride (15 mg/kg); additional anesthetic doses were administered as needed. Animals were then mounted in a stereotaxic apparatus, a small craniotomy was made over the right CgC, and the dura was reflected. A glass pipette loaded with rAAV-pEF1a-DIO-FLPo-WPRE-hGHpA (retrograde AAV; addgene, Cat. No. 105714) and AAV1-Syn-ChrimsonR-tdTomato (marker AAV; addgene, Cat. No. 105714) to mark the cortical injection site (ratio: marker AAV to rAAV = 1:2) was then slowly advanced into the right CgC using the coordinates (in mm): 1.0 caudal, 0.5 lateral, and 0.8 dorsal to the bregma. A total volume of 100 nl of AAV at a titer greater than 1×10^12^ was slowly infused. One week later (Figure 1C), mice were again subjected to the same surgical procedures for stereotactic infusion of 100 nl of AAV8-Ef1a-fDIO-GCaMP6s (addgene, Cat. No. 105714) at a titer greater than 1×10^13^ into the bilateral LC using the following coordinates (in mm): −5.4 caudal, ±0.8 lateral, and 3.0 dorsal to the bregma. Both sets of coordinates for the CgC and LC were determined according to the Allen Brain Atlas. The tip of the glass pipette was bevelled to an angle of ∼40° with a tip opening of 60-80 μm using a microelectrode beveler (BV-10, Sutter Instrument, Novato CA, USA) and connected to a 2.0 μL Hamilton syringe (Neuros Model 7002 KH, Hamilton, Reno, NV, USA). AAV was delivered manually at a rate of 5 nl/min. Mice were euthanized for detailed histological examination 10 weeks after the second AAV infusion into the LCs (Figure 1C).

### Perfusion, Sections and IHC

Mice were deeply anesthetized with urethane (1.3 g/kg) and perfused via the cardiovascular system with 4% paraformaldehyde in 0.1 M phosphate buffer (PB). The brain was dissected and postfixed in the same fixative overnight at 4°C. After fixation, the brain was blocked into forebrain and brainstem. The brainstem was cut into 50 μm thick sections in the coronal plane using a vibratome (Lieca VT1000 S, Germany), and serial sections containing the LC were collected and stained with antibodies against TH (Figure 1C). The experiment was continued only if TH antibody staining showed that the number of labeled LC-NE neurons in the bilateral LC was less than a single digit and all labeled neurons showed TH immunoreactivity (IR) (Figure 2). In this case, the forebrain block (from Bregma 1.2 to −3.7) was sectioned with a vibratome into 50 μm thick sections in the coronal plane, and every third section of the total consecutive forebrain sections was stained with IHC and examined (Figure 1C). Of these, those covering the CgC were stained by IHC with primary antibodies against TH and GCaMP6s (GFP) labeled with secondary antibodies conjugated to fluorophores with broad absorption/emission spectra (Figure 3). This fluorescence IHC (FIHC) staining was used to validate the viral infusion site showing tdTomato signal (marker AAV) and to confirm that GFP-IR fibers also showed TH-IR and thus originated from the LC. If any GFP-IR fibers were found that did not show TH-IR, the experiment was terminated. The remaining sections were stained with the primary antibodies against GFP alone, followed by staining with biotinylated secondary antibodies and avidin-biotin complex (ABC; VECTASTAIN® Elite® ABC-HRP kit; VectorLabs) (Figure 4A); immunoreactivity products were visualized using 3,3’-diaminobenzidine (DAB) as chromogen. After IHC, FIHC sections were stained with fluorescent Nissl; sections stained with IHC combined with the ABC method (IHC/ABC) and those not used for IHC were stained with conventional Nissl to aid facilitate identification of cortical regions.

**Figure 2.**
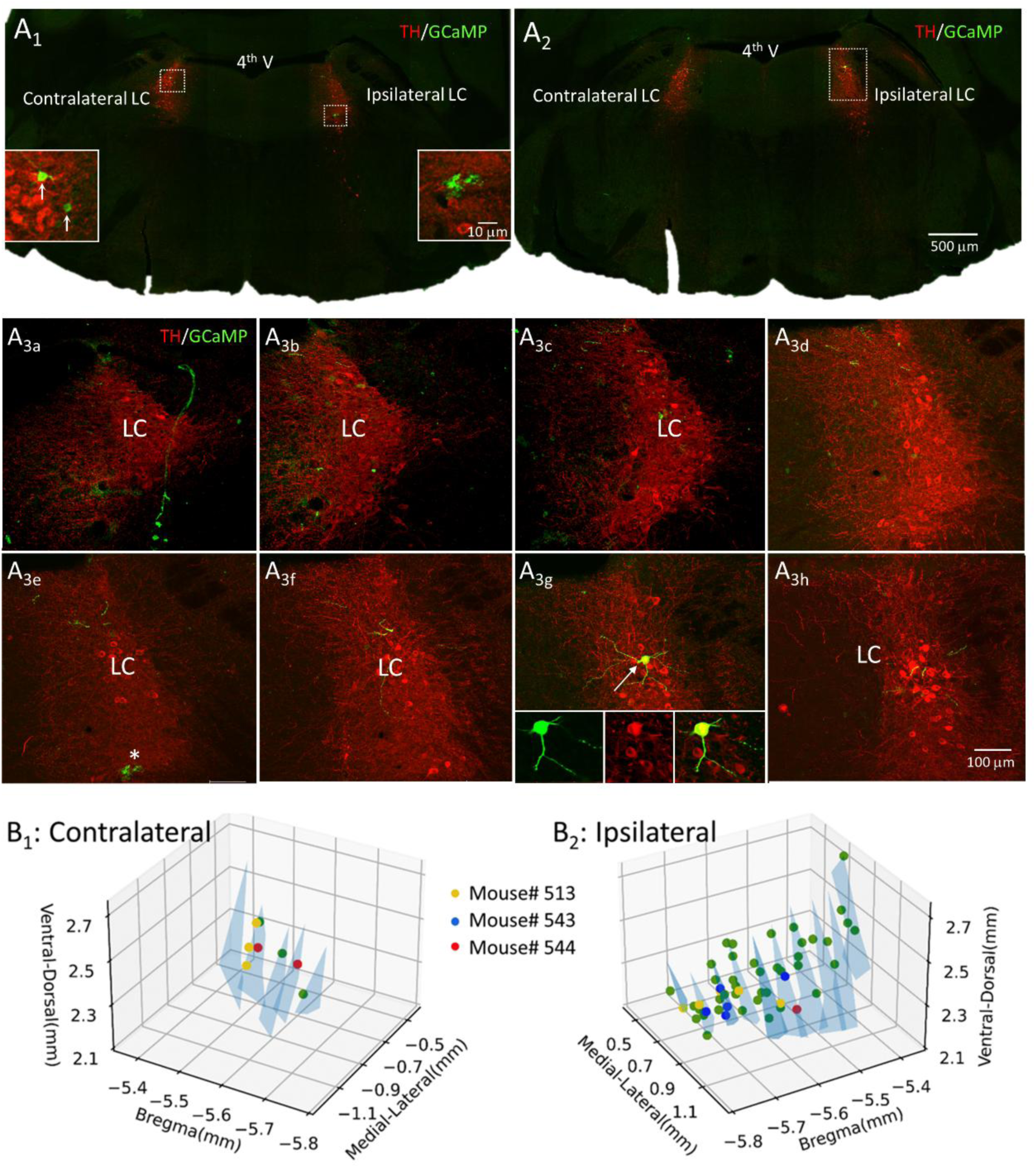
Representative result shows the labeling of NE neurons in the bilateral LC using the modified TrAC method. The data are from the study of mouse #544. ***A***. Low-power images show two pontine sections containing the LC that were stained with primary antibodies against TH (red) and GFP (green). Two neurons are labeled in the rostral LC section (A1) and one in the caudal section (A2). The two neurons in the rostral section (A1) are contralateral to the rAAV infusion, while the neuron in the caudal section (A2) is ipsilateral. The lower left inset of the rostral section (A1) is an enlargement of the dotted area of the contralateral LC, showing the two labeled neurons (arrows). The lower right inset shows that the structures in the dashed area of the ipsilateral side of the rostral section are not neurons (see also the asterisk in A3e). Eight high-power images (A3) show the entire ipsilateral LC. Image A3g shows the cell body and proximal dendrites of the labeled neuron and is from the same caudal section shown in the low-power (A2). The insets at the bottom (A3g) show the labeled neuron exhibiting TH-IR. ***B***. Plots show the distribution of all labeled NE (TH-and GFP-IR) neurons in the contralateral (B1) and ipsilateral (B2) LC, summarized from the five study cases. Yellow, blue, and red circles denote data from mice #513, #543, and #544, respectively; a green circle denotes data from the remaining two study cases. Note that no contralateral neurons were labeled in the study of mouse #543. The light blue triangular shadow delineates the LC.

**Figure 3.**
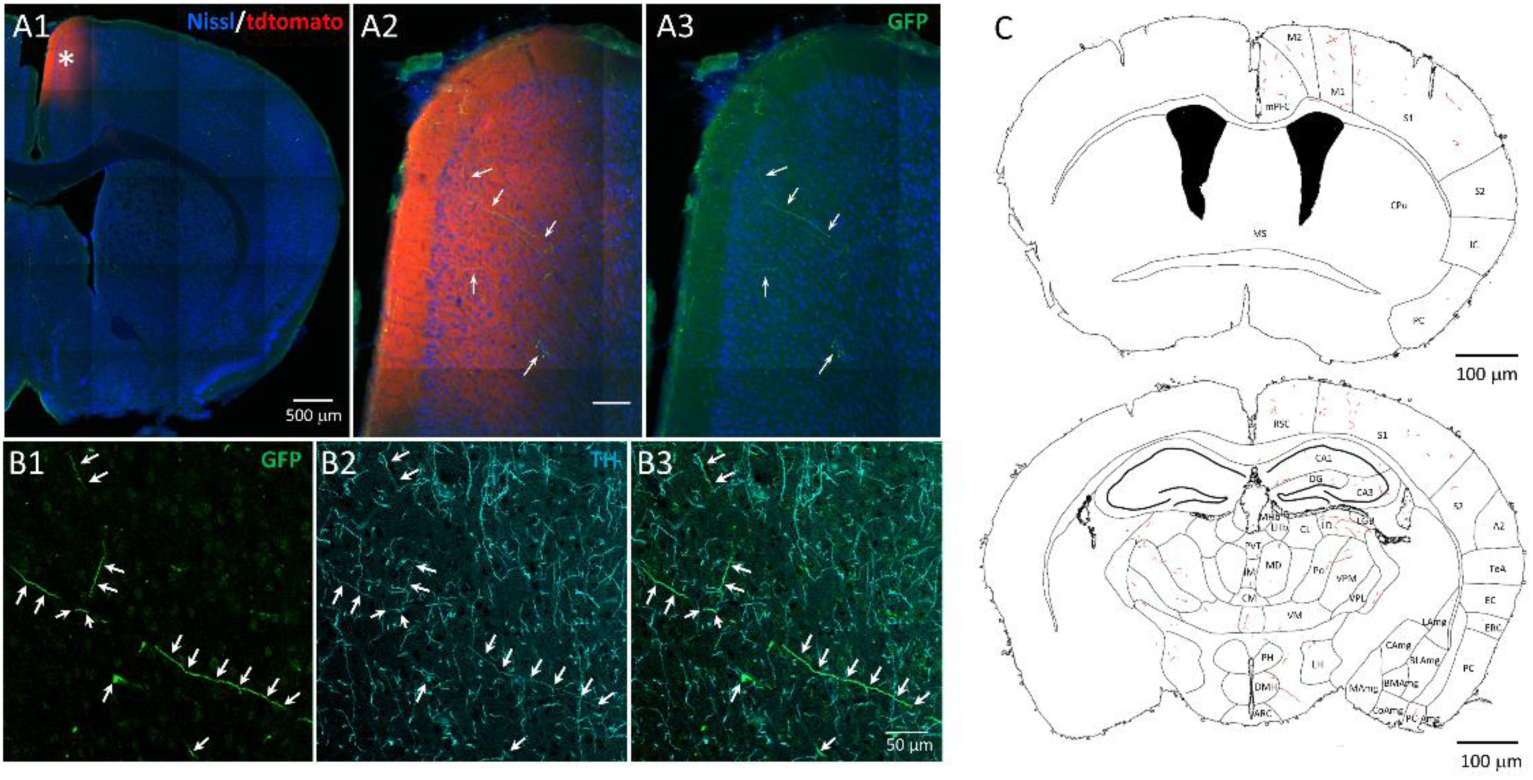
Representative result showing the tracing of NE (TH and GFP-IR) axons in the CgC using the modified TrAC method. The data are from mouse #543. **A**. A low-power merged image of tdTomato (red), GFP (green), and Nissl (blue) staining of a forebrain section shows the viral infusion site (marked by an asterisk) in the CgC. Magnification of the field reveals the GFP (GCaMP6s)-IR fibers (as indicated by the arrows in panel A2). These fibers are more visible when only the GFP signal is shown (Figure A3). **B**. Higher-power images show the same field from the next consecutive section, which was stained with anti-GFP antibodies (green in B1) and anti-TH antibodies (cyan in B2), as well as a merged image (B3). Note that all GFP-IR fibers also exhibit TH-IR, as indicated by the arrows. **C**. Representative plots show the complete tracking of NE fibers in two sections stained with IHC/ABC. Note the unilateral and bilateral distribution of NE fibers in the cortex and thalamus (bottom). *Brain region abbreviations*: **A2**: Secondary auditory cortex; **ARC**: arcuate area (of the hypothalamic area); **BLAmg**: Basolateral amygdala nucleus; **BMAmg**: Basomedial amygdala nucleus; **CA1**: Cornu ammonis area 1 (of the hippocampus); **CA3**: Cornu ammonis area 3 (of the hippocampus); **CAmg**: Central amygdala nucleus; **CL**: Centrolateral thalamic nucleus; **CM**: Centromedial thalamic nucleus; **CoAmg**: Cortical amygdala nucleus; **CPu**: Caudate putmen (striatum); **DG**: Dentate gyrus (of the hippocampus); **DMH**: Dorsomedial hypothalamic area; **EC**: Ectorhinal cortex; **ERC**: Entorhinal cortex; **IC**: Insular cortex; **IM**: Intermedial thalamic nucleus; **LAmg**: Lateral amygdala nucleus; **LD**: Laterodorsal thalamic nucleus; **LGB**: lateral geniculate body; **LH**: Lateral hypothalamic area; **LHb**: Lateral habenular nucleus; **M1**: Primary motor cortex; **M2**: Secondary motor Cortex; **MAmg**: Medial amygdala nucleus; **MD**: Mediodorsal thalamic nucleus; **MHb**: Medial habenular nucleus; **mPFC**: Medial prefrontal cortex; **MS**: Medial septal nucleus; **PC**: Piliform cortex; **PCAmg**: Piriform-amygdala area; **PH**: Posterior hypothalamic area; **Po**: Posterior thalamic nucleus; **PVT**: Paraventricular thalamic nucleus; **RSC**: Retrosplenial cortex; **S1**: Primary somatosensory cortex; **S2**: Secondary somatosensory cortex; **TeA**: Temporal association cortex; **VM**: Ventromedial thalamic nucleus; **VPL**: Ventral posterolateral thalamic nucleus; **VPM**: Ventral posteromedial thalamic nucleus.

**Figure 4.**
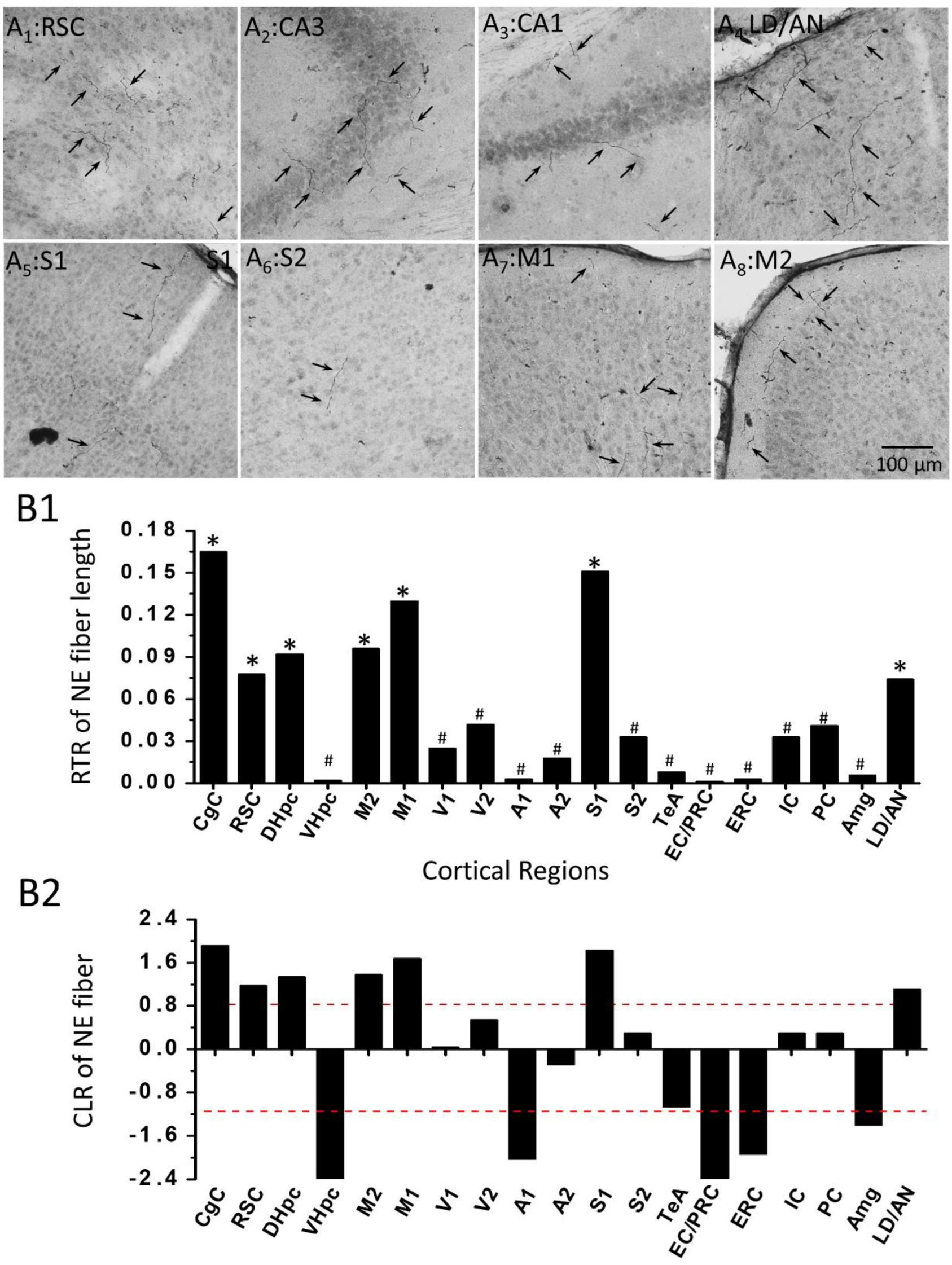
Representative images show NE axon tracing in cortical regions other than the CgC, using the modified TrAC and anti-GFP IHC-ABC methods. These data are from the study of mouse #543. **A**. Representative images of sections that do not cover the CgC (rAAV infusion site) and are stained with anti-GFP IHC-ABC. The images show NE fibers in the RSC (A1), CA3 (A2), CA1 (A3), DL/AN (A4), S1 (A5), S2 (A6), M1 (A7), and M2 (A8). **B**. The plot shows the RTR histogram of NE fiber length of mouse #543. * denotes PTs, and # denotes cortical and hippocampal regions have CLR values less than up-threshold. The red dotted lines show up- and low-thresholds of CLR values. Cortical region abbreviations: **A1**: Primary auditory cortex; **A2**: Secondary auditory cortex; **Amg**: Amygdala; **AN**: anterior nucleus (of the thalamus); **CgC**: cingulate cortex; **DHpc**: Dorsal hippocampus; **EC/PRC**: Ectorhinal cortex/ perirhinal cortex; **ERC**, entorhinal cortex; **IC**: Insular cortex; **LD**: laterodorsal nucleus (of the thalamus); **M1**: Primary motor cortex; **M2**: secondary motor cortex; **PC**: Pontine cortex; **RSC**: retrosplenial cortex; **S1**: Primary somatosensory cortex; **S2**: Secondary somatosensory cortex; **TeA**: Temporal association area; **V1**: Primary visual cortex; **V2**: Secondary visual cortex; **VHpc**: Ventral hippocampus.

For FIHC, sections were rinsed briefly with PB followed by phosphate-buffered saline (PBS; 0.9% NaCl in PB). Sections were incubated for 1 hour with a solution consisting of 0.1% bovine serum albumin (BSA) in PBST (0.3% Triton in PBS) to block nonspecific reactions. Sections were then incubated overnight at 4°C in a primary antibody cocktail including rabbit anti-TH (1:1000 dilution; Cat. No. 6A2907, Merck Millipore, Burlington, MA, USA) and chicken anti-GFP (1:2000 dilution; Cat. No. ab13970, Abcam Inc., Cambridge, UK). For sections covering the CgC, the tdTomato (ChrisomR-tdTomato) signal was detected directly without further antibody staining. Sections were then incubated with a secondary antibody cocktail including donkey anti-rabbit IgG conjugated to Alexa FluorTM Plus −594 (1:600 dilutions; Cat. No. A-32754, Thermo Fisher Scientific, Waltham, MA, USA) and goat anti-chicken IgY conjugated to CF®488A (1:500 dilution; Cat. No. Biotium#20020, Biotium, Fremont, CA, USA) in PBST for 2 hours. Sections were washed three times for 10 minutes with PBST between antibody incubations. For the IHC/ABC method, primary antibody incubation was as described for FIHC, except that sections were first treated with 3% H_2_O_2_ in 0.1 M PB for 20 minutes to quench endogenous peroxidase activity, followed by rinsing in 0.1 M PB and PBST, and the same chicken anti-GFP antibodies were used alone. After washing in PBST, sections were incubated with biotinylated goat anti-chicken IgY secondary antibody (1:200 dilution; Cat. No. BA-9010; VectorLabs, Burlingame, CA, USA) in PBST for 1 hour, followed by washing in PBST and incubation with avidin-biotin-horseradish peroxidase complex (VECTASTAIN® Elite® ABC-HRP Kit; Vector Labs) for 1-2 hours. After five washes with PBS, sections were immersed in 0.0125% DAB solution for 10 minutes, 0.002% H_2_O_2_ was added, and sections were incubated for an additional 10 minutes.

### Data Analysis

After completion of the staining procedures, sections containing CgC were examined and photographed using a Zeiss LSM 780 confocal microscope system (Carl Zeiss); the other sections were examined and photographed using a light microscope system (Zeiss axio observer Z1 with Zen blue 3.1 software). For each section, the entire section was photographed. The NE fibers and the outlines and landmarks of the section were traced using ImageJ software. The cortical areas constituting the section were identified by the Nissl method using QuickNII software. After axon tracing was completed in all examined forebrain sections, the lengths of the traced NE fibers were measured. Then, a histogram of the summed NE fiber length relative to the total fiber length in the cortex (region-to-total ratio, RTR) was plotted for each cortical region. To access the relatively preferential cortical targets of the LC-NE retrogradely labeled with rAAV infused into the CGC, the CLR transformation of RTR was performed using the following equation (Aitchison, 1986):

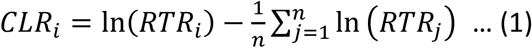

where CLR_i_ and RTR_i_ are the CLR and RTR of a given cortical region, respectively, and n is the number of cortical regions where NE fibers are found. For practical CLR calculations, equation 1 is written as follows:

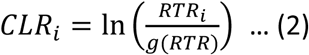

where *g(RTR)* represents the geometric mean of all components, which can be calculated as

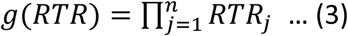

To identify relatively PTAs of the LC-NE neurons retrogradely labeled with rAAV infused into the CgC, a CLR threshold was obtained that could objectively indicate enrichment relative to the geometric mean. Namely, the CLRs of all the study cases were pooled and clustered using the K-means algorithm (Lloyd, 1982), which partitions data into clusters by minimizing the within-cluster sum of squares (WCSS). The elbow method was also used to identify additional clusters providing diminishing returns in WCSS reduction, and silhouette analysis was used to evaluate cluster cohesion and separation.

## Results

### CgC-projecting NE neurons appear to be randomly distributed in the LC

After a series of proof-of-concept experiments, we found that the time interval of 1 week between the infusion of rAAV (together with the marker AAV for the expression of tDtomato to mark the cortical injection site) into the CgC and AAV8 for the Flp-dependent expression of GCaMP6s into the LC, as well as the optimal survival time of 10 weeks after AAV8 infusion, allowed the best results. Notably, we found no NE fibers in the cortex with a survival time of less than 8 weeks after AAV8 infusion, although neurons with TH-IR were observed in the LC. Accordingly, we performed 10 experiments using the protocol described above (Figure 1C). Examination of LC sections stained with anti-TH antibody revealed that most of the GCaMP6s (GFP)-positive neurons also exhibited TH-IR in the LC, hereafter referred to as NE neurons, in 5 cases. The other 5 cases failed for unknown reasons because no NE neurons were found in the LC. Of the 5 successful cases, 40 NE neurons were labeled in one case, 5 NE neurons plus one neuron showing no TH-IR were labeled in one case, 5 NE neurons were labeled in two cases, and even better, a single NE neuron was labeled in one case (Figure 2A) in the LC ipsilateral to the cortical injection site. When the location of the 61 NE neurons from the 5 cases was constructed with registration according to the 4th ventricle using QuickNII software, it appears that these projection neurons to the CgC were randomly distributed rather than clustered at a specific location in the LC (Figure 2B). This observation suggests that NE neurons projecting to the CgC do not have a specific topographical location in the LC, an observation consistent with that of Uematsu et al. (2017). Notably, there were also labeled LC-NE neurons in the contralateral LC; the total number of labeled neurons from the 5 cases is 8 cells, and again they appear to be randomly distributed in the contralateral LC (Figure 2B).

### LC-NE neurons projecting to the mPFC show a feature of lateralized projection

We further quantitatively analyzed forebrain sections covering the CgC of the 3 cases, designated Mouse# 513, Mouse# 543, and Mouse# 544, in which only 5 (2 cases) and 1 NE neuron (1 case) were identified in the ipsilateral LC. The remaining 2 cases were not further investigated due to too many labeled cells (40) and the presence of a cell that did not show TH-IR in the ipsilateral LC. We confirmed that the examined GFP-IR fibers at the infusion site showed TH-IR (Figure 3A, B), hence GFP-IR, hereafter referred to as NE fibers. Interestingly, NE fibers were mostly found in the cortical areas and hippocampus ipsilateral to the injection side, while very rarely in the contralateral side (Figure 3C; Supplementary Figures 1-3). This observation suggests that although unilateral CgC infusion of rAAV labeled NE neurons on both sides of the LC, these labeled LC-NE neurons appear to have a lateralized projection to the cortex and hippocampus; a similar result has been reported for LC-NE neurons projecting to the spinal cord (Hirschberg et al. 2017). Accordingly, we focused on a quantitative analysis of NE fiber distribution in the right cortex.

### Axon projection from sparse LC-NE neurons to the cortex is divergent but not uniform

After axon tracing was completed in all examined forebrain sections, the lengths of the traced NE fibers were measured and a RTR histogram of NE fiber length was plotted. Interestingly, the results showed that a small number of LC-NE neurons projected NE fibers divergently across many cortical regions rather than being restricted to a few regions (see Figures 3C and 4, as well as Supplementary Figures 1-3). However, this pattern is not uniform. There are a few cortical regions with an RTR greater than or approximately equal to CgC (see the asterisk in Figure 4B1), as well as more regions with much lower RTRs (see the octothorpe in Figure 4B1). To gain insight into the RTR histogram of NE fiber length across cortical and hippocampal regions, a CLR transformation was applied to normalize the RTR. This analysis revealed several PTAs with marked enrichment of NE fiber length in these areas relative to the geometric mean of all other areas. The K-means algorithm, together with elbow and silhouette analyses, determined an optimal cluster number of K = 3, separated by up-and low-thresholds (CLR = 0.83 and −1.14, respectively). These three clusters represent regions that are enriched (CLR > 0.83), proportional (−1.14 < CLR < 0.08), or depleted (CLR < −1.14) relative to the geometric mean. Theoretically, rAAV labeled LC-NE neurons because they should have a relatively large number of axonal projections to the CgC. Supporting this argument, we found that the CgC had a CLR value much higher than the up-threshold in both Mouse #543 and Mouse #544. In Mouse #513, however, the CgC had a CLR value of 0.4. This study case had more labeled LC-NE neurons, as well as more cortical and hippocampal regions where NE fibers were found. Therefore, it is likely that more brain regions with NE fibers resulted in apportionment of the NE fibers, thereby lowering the RTR and CLR of the CgC. Based on this analytical strategy, we considered cortical and hippocampal regions with a CLR value higher than the up-threshold to be the PTAs of the labeled LC-NE neurons’ axonal projections. The following is a case-by-case description of the functional implications of the labeled LC-NE neuron projections to the cortex and hippocampus according to this analytical strategy.

### The PTs of axonal projections from sparse LC-NE neurons are functionally correlated

Figure 4B1 shows the RTR histogram of NE fiber length for mouse #543. The core rAAV injection site is at approximately Bregma 0.66 and covers mostly the CgC of the mPFC, as well as a smaller portion of motor cortex 2 (M2) (see the asterisk in Figure 3A1 and the yellow shadow in Supplemental Figure 1). Five NE neurons were labeled in the ipsilateral LC and none in the contralateral LC (see the blue circle in Figure 2B). Despite the low total number of labeled neurons (five cells), the NE fiber distribution in the cortex is divergent but not uniform, as can be seen. The CLRs in the CgC, S1, M1/M2, retrosplenial cortex (RSC), and dorsal hippocampal cortex were higher than the up-threshold. These regions are considered PTAs of the five LC-NE neurons (Figure 4B2). Importantly, previous studies have shown functional correlation of the defined TPAs (the CgC, RSC, and DHpc) for spatial cognition (Vann et al., 2009; Mitchell et al., 2018). In contrast, the CLR in the ventral hippocampal cortex (VHpc) is much lower than the low threshold.

In mouse #544, the core cortical injection site was predominantly located in the CgC (approximately at Bregma 0.98; see the yellow shadow in Supplemental Figure 2). One NE neuron was labeled in the ipsilateral LC, and two were labeled in the contralateral LC (red circle in Figure 2B). Although fewer NE neurons were labeled than in mouse #543, the distribution of NE fibers across the cortex and hippocampus diverged again (Figure 5A). In addition to the CgC, the S1, RSC, and DHpc are classified as TPAs of LC-NE neuron projections because their CLR values are higher than the up-threshold (Figure 5B). These TPAs resemble those of mouse #543 except that no motor region is included, reflecting the lower number of labeled NE neurons.

**Figure 5.**
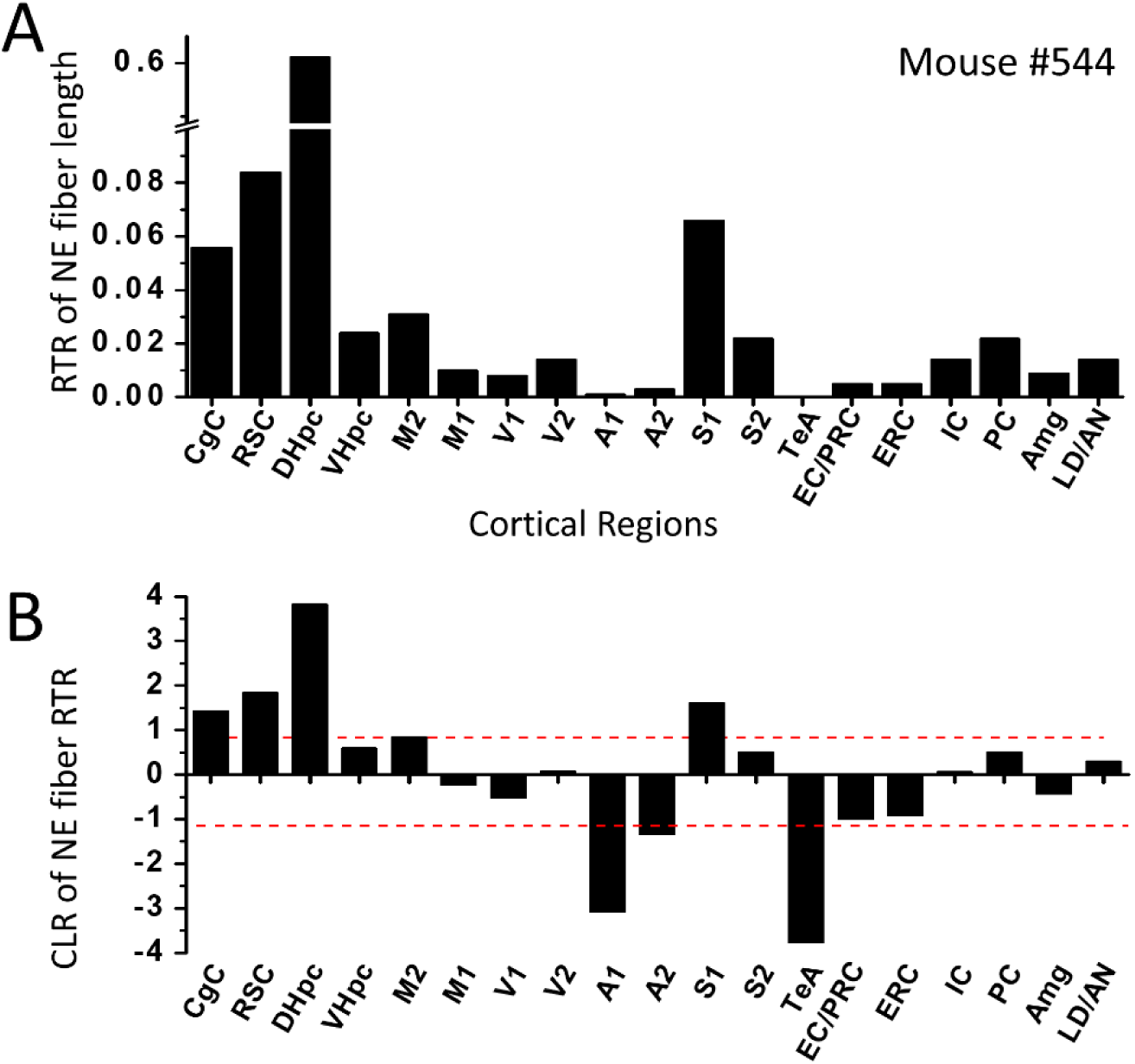
The plots show RTR histograms of NE fiber length (A) and the CLR (B) of mouse #544. The red dotted lines show the up- and low-thresholds of the CLR. See Figure 4 for abbreviations of cortical regions.

In mouse #513, the core cortical injection site was located in the anterior CgC (approximately at Bregma 0.26), extending into the M2 cortex (see the yellow shadow in Supplemental Figure 3). In this case, five NE neurons were labeled in the ipsilateral LC and three in the contralateral LC (see the yellow circle in Figure 2B). Once again, even when the total number of labeled NE neurons was as few as eight, the NE fiber distribution across the cortex diverged (Figure 6A). The DHpc, S1/S2, and the olfactory piriform cortex (PC) had CLR values higher than the up-threshold (Figure 6B). Additionally, the CgC, RSC, M2, and VHpc had CLR values higher than zero but lower than the up-threshold (Figure 6B). All other cortical regions had negative CLR values except extrastriate visual area 2 (V2), which had a low CLR value of 0.069 (Figure 6B). Thus, this study is consistent with the previous two in that the defined TPAs are functionally correlated for spatial navigation, even though the CgC and RSC had CLR values below the up-threshold. Still, these values are much higher than those of the other regions. Compared to the previous two, this study case had more cortical regions that were defined as TPAs or had CLR values approaching the up-threshold, possibly due to a greater number of labeled LC-NE neurons. Interestingly, in addition to CgC, RSC, and DHpc, these regions were sensory (S1, S2, V2, and PC) or motor (M1 and M2). Unlike the previous two, the VHpc had a CLR value similar to those of the CgC and RSC in mouse #513,.

**Figure 6.**
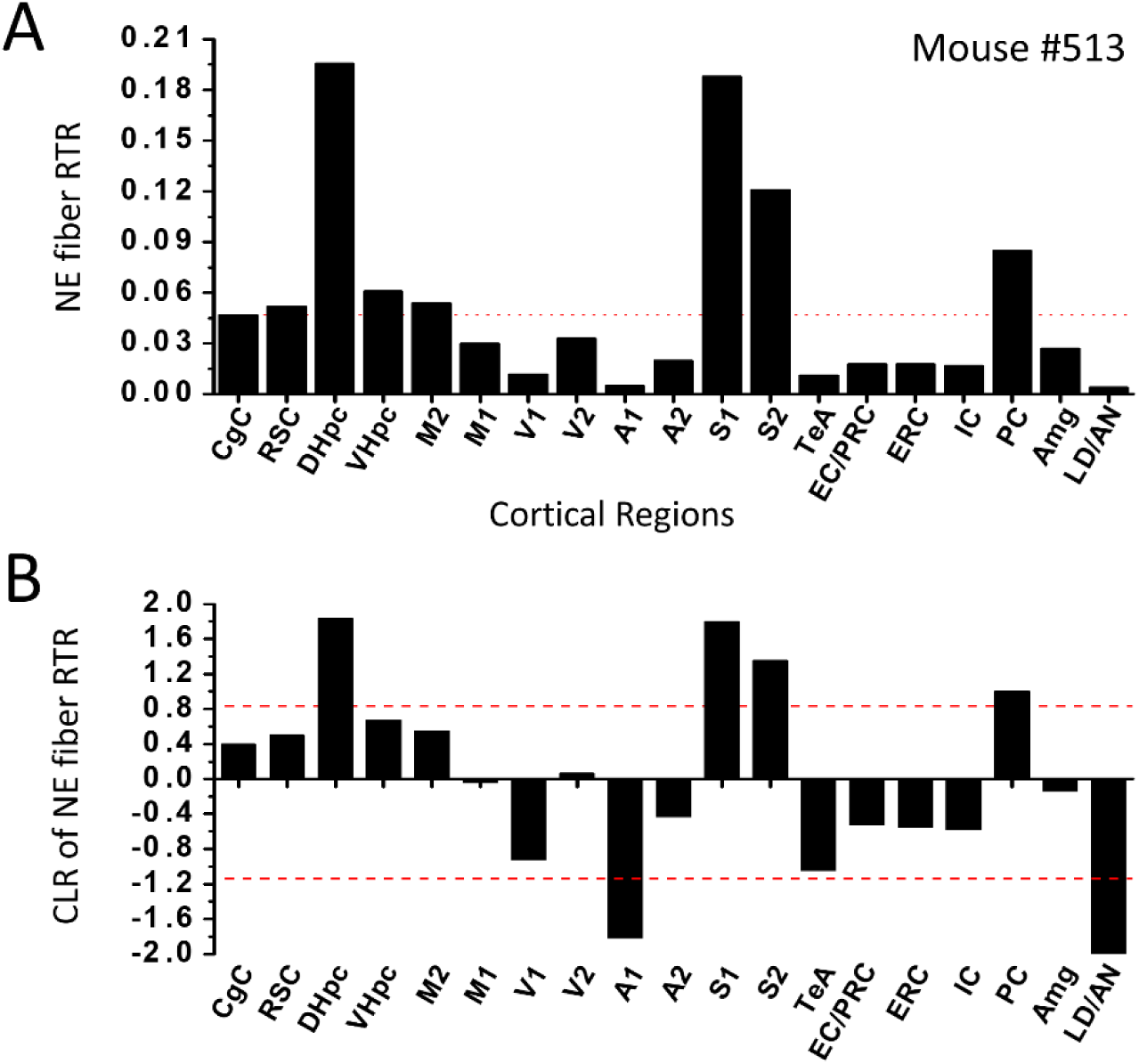
The plots show RTR histograms of NE fiber length (A) and the CLR (B) of mouse. The red dotted lines show the up- and low-thresholds of the CLR. See Figure 4 for abbreviations of cortical regions.

In summary, the results consistently define the CgC, RSC, and DHpc as TPAs of axonal projections from spare LC-NE neurons. Depending on the number of labeled LC-NE neurons, some sensory and motor regions may also be defined as TPAs. The more LC-NE neurons labeled, the more sensory and motor regions can be defined as TPAs. This feature reveals that injecting rAAV into the CgC activates a small number of LC-NE neurons with extensive axonal projections to cortical regions involved in high-level cognitive functions such as decision-making, spatial memory, navigation, and sensory and motor functions. Therefore, this small number of LC-NE neurons appears capable of orchestrating goal-directed navigation by coordinating neuronal activity in their TPs. This argument is further supported by averaging the RTR histogram of the three study cases (Figure 7). Since the cortical regions CgC, RSC, and DHpc, as well as the anterior and laterodorsal nuclei of the thalamus, are key structures in the navigation network (Vann et al., 2009; Mitchell et al., 2018), we also analyzed the RTRs in these two thalamic nuclei. The results show that labeled LC-NE neurons generally have divergent projections to the thalamus (Figure 3C and Supplementary Figures 1–3). Interestingly, unlike the cortex and hippocampus, the projections appear to be bilateral in mice #543 and #544 (see Supplementary Figures 1 and 2). However, the RTRs of the LD/AN do not meet the criteria of any of the TPAs, except in mouse #543 (Figure 4B2).

**Figure 7.**
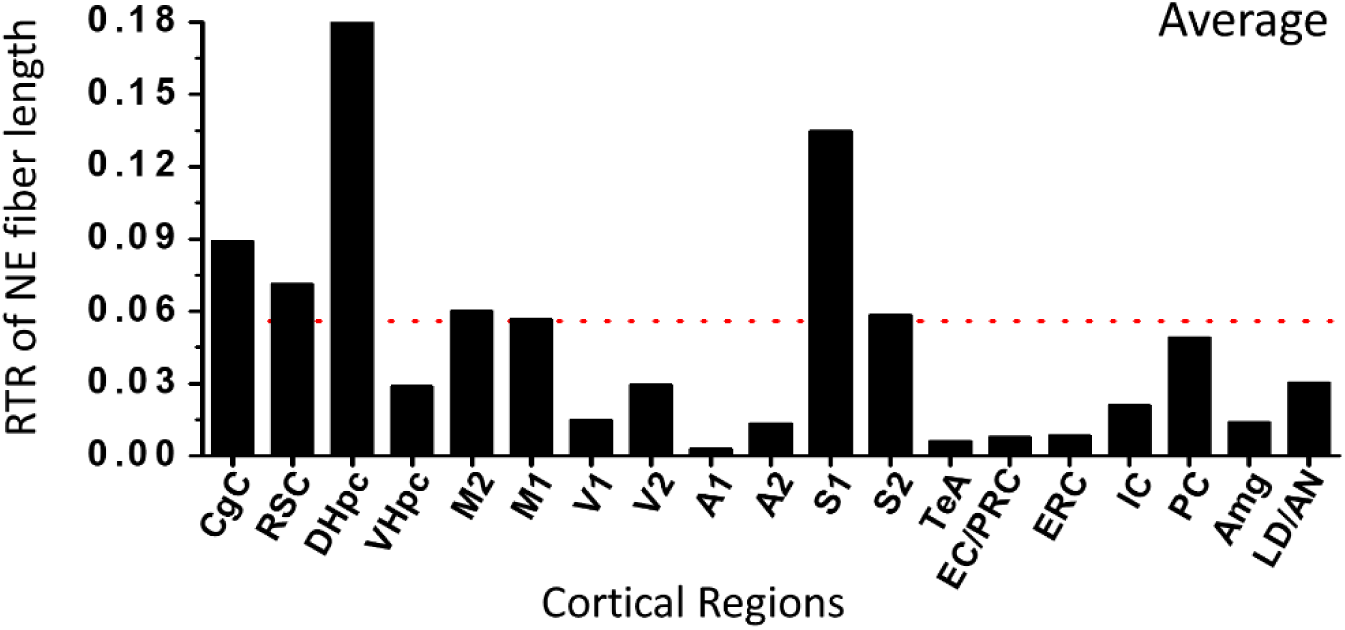
The plot show RTR histogram of NE fiber length of the three study cases. See Figure 4 for abbreviations of cortical regions.

## Discussion

### Technical considerations

In this study, the TrAC protocol of Plummer et al. (2020) was modified to sparsely label LC-NE neurons, taking advantage of the fact that viral vectors typically do not achieve complete transfection. We hypothesized that injection of rAAV would selectively label a subset of LC-NE neurons projecting to the CgC, followed by a second AAV for Flp-dependent GCaMP6s expression, further limiting the number of labeled neurons. Our approach resulted in a 30% success rate (3 out of 10 experiments) in labeling fewer than 10 neurons, which is a significant step toward better visualization and understanding of LC-NE neuronal projections. GCaMP6s was chosen as the FT not only because of its superior axonal transport properties, but also to test the possibility that the method achieved for high specificity and sparse labeling of LC-NE neurons could be applied to future studies aimed at high-resolution imaging of neuronal activity coordinated by the LC-NE system. The use of a dual recombinase system ensured specificity, as GFP-IR fibers observed at the rAAV injection site also exhibited TH-IR. Since there is currently no evidence that neurons in the LC express only TH and not DBH, and our previous study has shown that our protocol for AAV injection into the LC is highly specific (Kuo et al., 2020), the observation that GFP-IR fibers are also TH-IR confirms their identity as noradrenergic fibers, and the possibility of staining NE and/or dopamine axons from sources other than the LC is low. However, it is possible that the NE (GFP-IR) fibers traced in this study also release dopamine, as the functional role of dopamine released from the LC has recently been firmly established (Takeuchi et al., 2016; Beas et al., 2018; Chowdhury et al., 2022). In summary, our approach reported here provides a cost-effective and accessible strategy using publicly available materials to advance the study of LC-NE and other neurons of the neuromodulatory system in behavioral and cognitive neuroscience.

### A single LC-NE neuron can influence multiple interconnected brain regions (TPAs)

Targeting the CgC for LC-NE labeling provides insight into how these LC-NE neurons orchestrate higher-order cognitive functions involving the CgC. Due to the low transfection efficiency of AAV, we believe that the NE neurons in the LC were retrogradely labeled by rAAV because they likely have relatively higher axonal projections to the CgC. Accordingly, the labeled LC-NE neurons should be able to effectively coordinate the function of the CgC and brain areas (PTAs) with similar or higher NE fiber innervation densities for adaptive behavior when phasic activation of the labeled LC-NE neurons is triggered by salient stimuli (Aston-Jones & Cohen, 2005). Our results also suggest that a single LC-NE neuron can have multiple brain regions as its PTAs, including the CgC and one or more others. Importantly, evidence from previous studies show that PTAs defined in all three study cases are functionally interconnected either directly or indirectly, and the function of the network is well established. Especially, the CgC, RSC, and DHpc are consistently defined as the PTAs in all three study cases, and their functions well correspond to neural networks involved in regulating attention, spatial navigation, and decision-making (Van Groen & Wyss, 1990, 1992, 2003; Jones et al., 2005; Kobayashi & Amaral, 2007; Vann et al., 2009; Mitchell et al., 2018). For the convenience of the further discussion, these three regions are referred to as core PTAs.

In the case of mouse #544, the number of labeled neurons is the sparsest (one ipsilateral neuron and two contralateral neurons). The NE fiber tracing results showed the S1, CgC, RSC, and DHpc to be the MjCPs. The S1’s functional relevance to the core MjCPs is evident, given that the RSC relays sensory input to the hippocampus (Vogt & Miller, 1983; Vann et al., 2009; Alexander et al., 2023). Together, by coordinating the neuronal activity in their PTAS, we speculate that the labeled LC-NE neurons in mouse #544 appear to modulate processes involved in decision-making requiring spatial memory and somatosensory cues. The study of mouse #543 revealed five ipsilateral and no contralateral labeled LC-NE neurons. This case study had a similar PTAs profile to that of mouse #544, with M1 and M2 being the additional defined PTAs. The functional correlation is evident here as well, as decision-making is typically followed by an adaptive response whose processes are regulated by the LC-NE system (Usher et al., 1999; Aston-Jones & Cohen, 2005), including the enhancement of sensorimotor gating (Alsene & Bakshi, 2011; Kuo et al., 2020). Taken together, the results of these two studies suggest that the more LC-NE neurons projecting to the CgC that are labeled, the more PTAs with functional correlations can be identified. This feature is also supported by the study of mouse #513. This study showed the largest number of labeled LC-NE neurons (five ipsilateral and three contralateral) and defined TPAs that largely overlap with the other two cases. As expected, additional sensory areas, such as the PC, S2, and V2, were also defined as PTAs. Interestingly, the VHpc was also identified as a PTA, indicating the presence of an LC-to-CgC/VHpc module capable of regulating more complex behaviors with emotional contexts (Strange et al., 2014). In summary, our axon tracing from spare neurons in the cortex demonstrates that the organization of cortical projection of a single LC-NE neuron allows the neuron to coordinate neuronal activity in multiple interconnected cortical regions. This feature of the cortical projection enables a limited number of LC-NE neurons to regulate complex brain functions and behaviors through different combinations of activated LC-NE neurons with different PTAs. It also supports the modular organization of LC-to-cortex projections. We hypothesize that the larger the number of activated LC-NE neurons, the larger the scale of neurocircuitry coordinated to optimize cognitive performance in complex environments. This also implies that cortical and subcortical inputs to the LC should be highly selective, activating appropriate LC-to-cortex projection modules for behavior regulation.

We also analyzed RTR in LD/AN of the thalamus because the interconnected network of the core PTAs that supports spatial memory and cognitive mapping (Van Groen & Wyss, 1990, 1992, 2003; Jones et al., 2005; Kobayashi & Amaral, 2007) also includes the LD/AN. Although the LD/AN did not consistently meet PTA criteria, it has the highest average RTR among PTAs (Figure 5C). We speculate that the labeled LC-NE neurons may still able to effectively regulate LD/AN neuron activity to interact with the core PTAs and the others due to the LD/AN’s high NE density, likely caused by their small structures compared to the CgC, RSC, and DHpc. Additionally, our data suggest that contralateral LC-NE neurons predominantly project contralaterally while ipsilateral neurons project ipsilaterally and may decussate at the thalamic level.

### All LC-NA neurons contribute to the maintenance of consciousness via non-PTAs

In addition to the PTAS, we were able to identify larger portions of the cortical projections of labeled LC-NE neurons as non-PTAs. Unlike PTAs, none-PTAS were more widely distributed, yet they had lower RTRs. Due to the low RTRs, it is unlikely that an LC-NE neuron would be labeled by rAAV infused into the CgC, which is one of the none-PTAs. In other words, it is unlikely that any of the labeled LC-NE neurons in this study contain only none-PTAs. Instead, we believe that the RTR pattern in each study case arises from the summation of labeled LC-NE neurons with different PTAs and none-PTAs. It is reasonable to deduce that such a cortical projection would allow LC-NE neurons to coordinate specific brain functions through their interconnected PTAs, while simultaneously contributing to global arousal regulation via their none-PTAs. Despite the low fiber density of none-PTAs in an individual LC-NE neuron, the collective none-PTAs of all LC-NE neurons may provide sufficient NE levels for arousal maintenance. Electron microscopy studies have suggested that NE exerts its effects through both synaptic and volumetric transmission mechanisms (Vizi et al., 2004; Min et al., 2007; Fuxe et al., 2015). We propose that, because it allows for high specificity in communicating with postsynaptic neurons, synaptic transmission would be more suitable for LC-NE neurons to modulate neuronal activity within the PTAs. In contrast, volumetric transmission supports a more diffuse and persistent influence across none-PTAs, maintaining cognitive vigilance and attentional readiness. Future studies should explore the relative contributions of these transmission modalities of LC-NE fibers across different cortical projection targets to test this speculation.

### Conclusions

Our study provides new insights into the cortical projections of LC-NE neurons labeled by rAAV infusion into the CgC. Key findings include: 1. CgC-projecting LC-NE neurons are randomly distributed throughout the LC, with no clear spatial clustering pattern. 2. The unilateral LC projects to both hemispheres, but individual LC-NE neurons exhibit lateralized, non-bidirectional projections. 3. Individual LC-NE neurons exhibit divergent, non-uniform projection patterns, characterized by dense PTAs and sparse none-PTAs. 4. PTAs are functionally interconnected and contribute to specific cognitive and behavioral processes. Specifically, the core PTAs identified in this study are known to form a neuronal network for spatial memory, navigation, and attentional control. These results suggest that individual LC-NE neurons serve as functional hubs that coordinate multiple cortical regions for specific behaviors. Meanwhile, all LC-NE neurons collectively contribute to the regulation of arousal through their widespread none-PTAs projections. This modular organization within the LC provides a flexible mechanism for coordinating neural networks, enabling adaptive behavioral responses in complex environments. Further research is needed to determine how different LC-NE neuron subgroup interact with different neural circuits, as well as how these interactions shape cognition and behavior. One way to achieve this would be to reveal more single or sparse LC-NE neurons with rAVV infused into different cortical regions.

## Acknowledgements

We thank the Technology Commons, College of Life Science, National Taiwan University for technical support.

