## Supplemental Figure 1 for "Axon Collateral Pattern of a Sparse Locus Coeruleus Norepinephrine Neuron in Mouse Cerebral Cortex"

### NE fibers (red) in forebrain sections of **Mouse #543**.

A: Bregma 1.26 mm

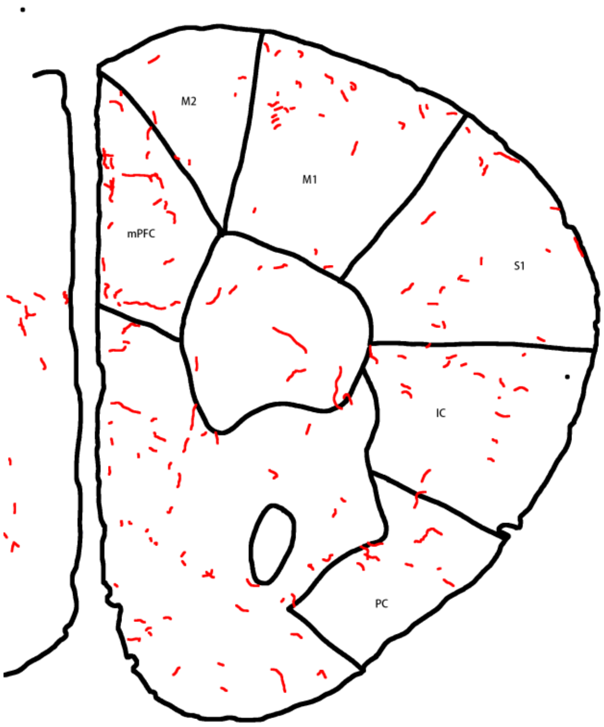


B: Bregma 0.79 mm
(Core injection site)

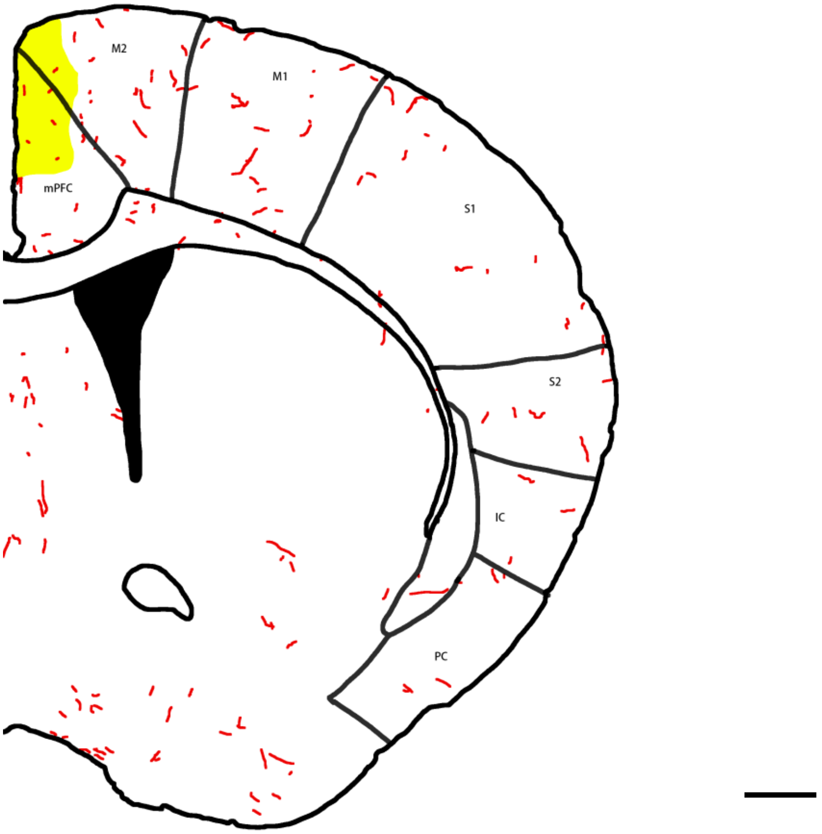

C: Bregma 0.66 mm

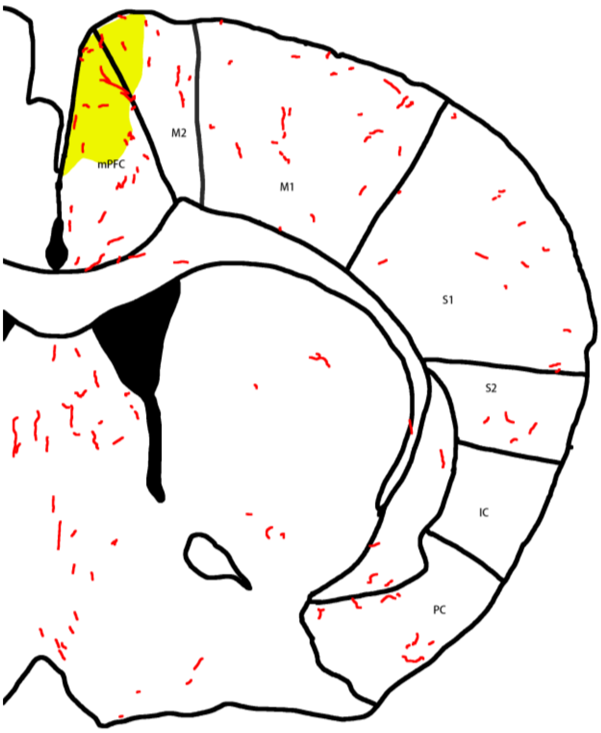


D: Bregma 0.58 mm

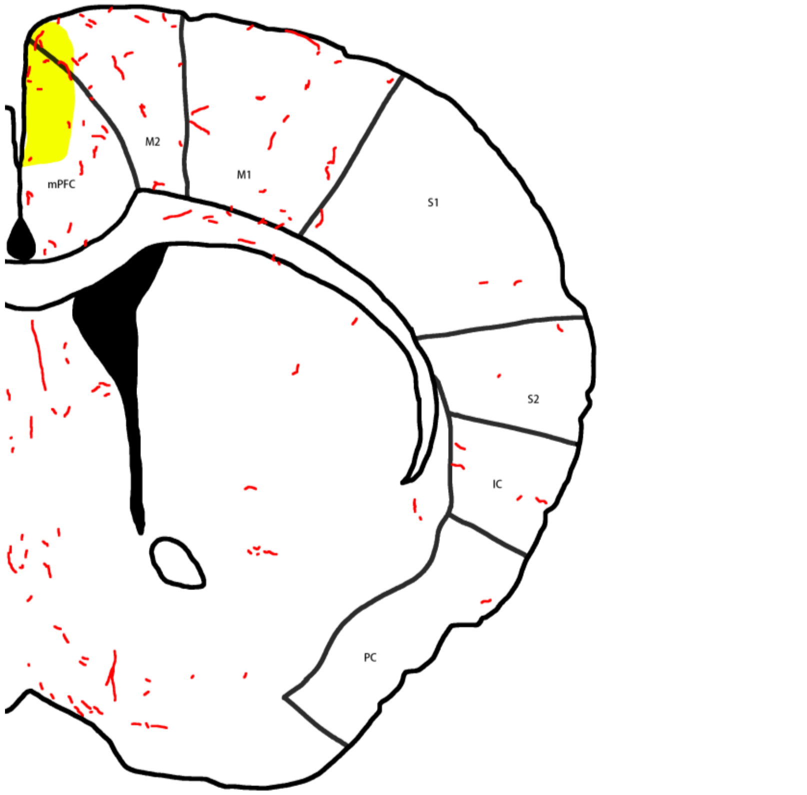

E: Bregma 0.43 mm

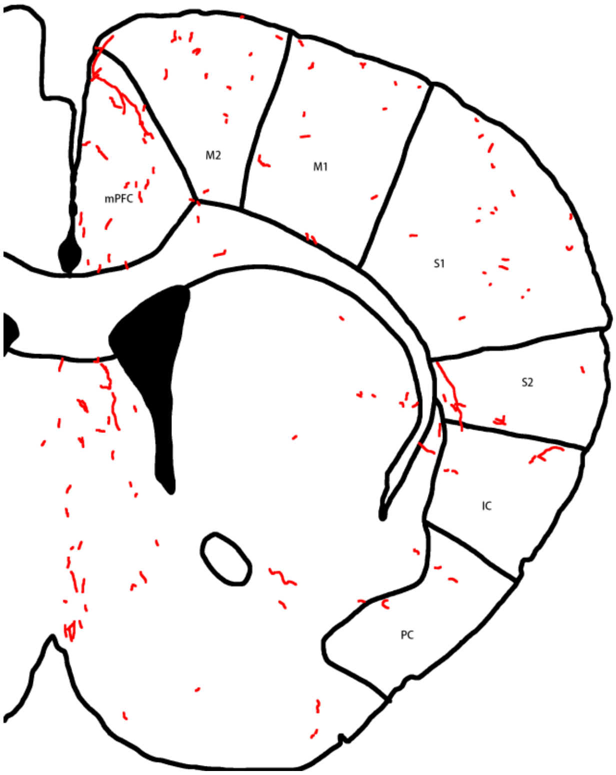


F: Bregma 0.31 mm

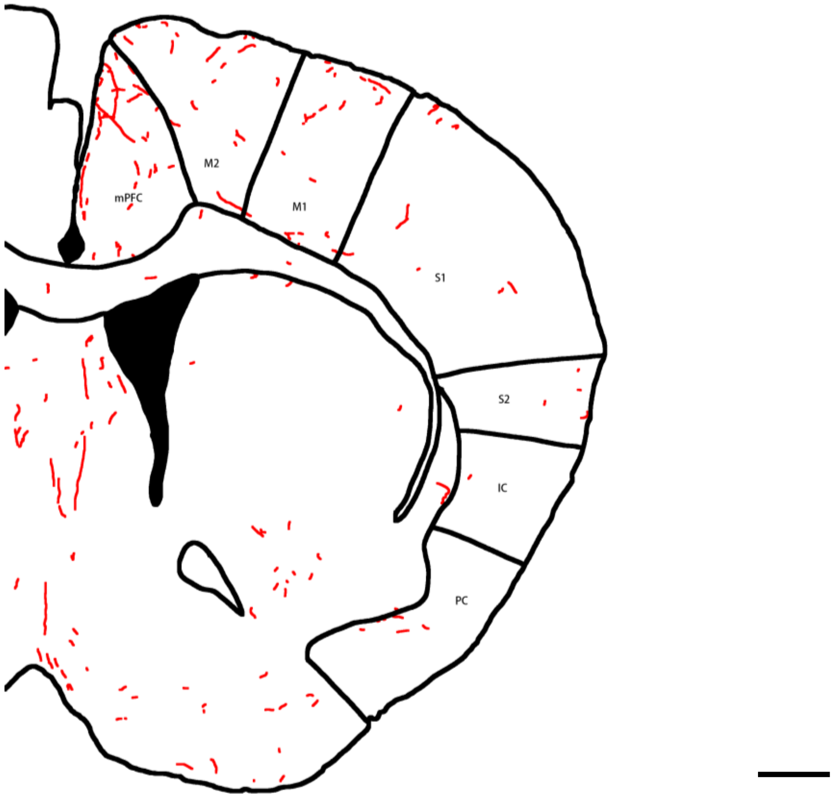

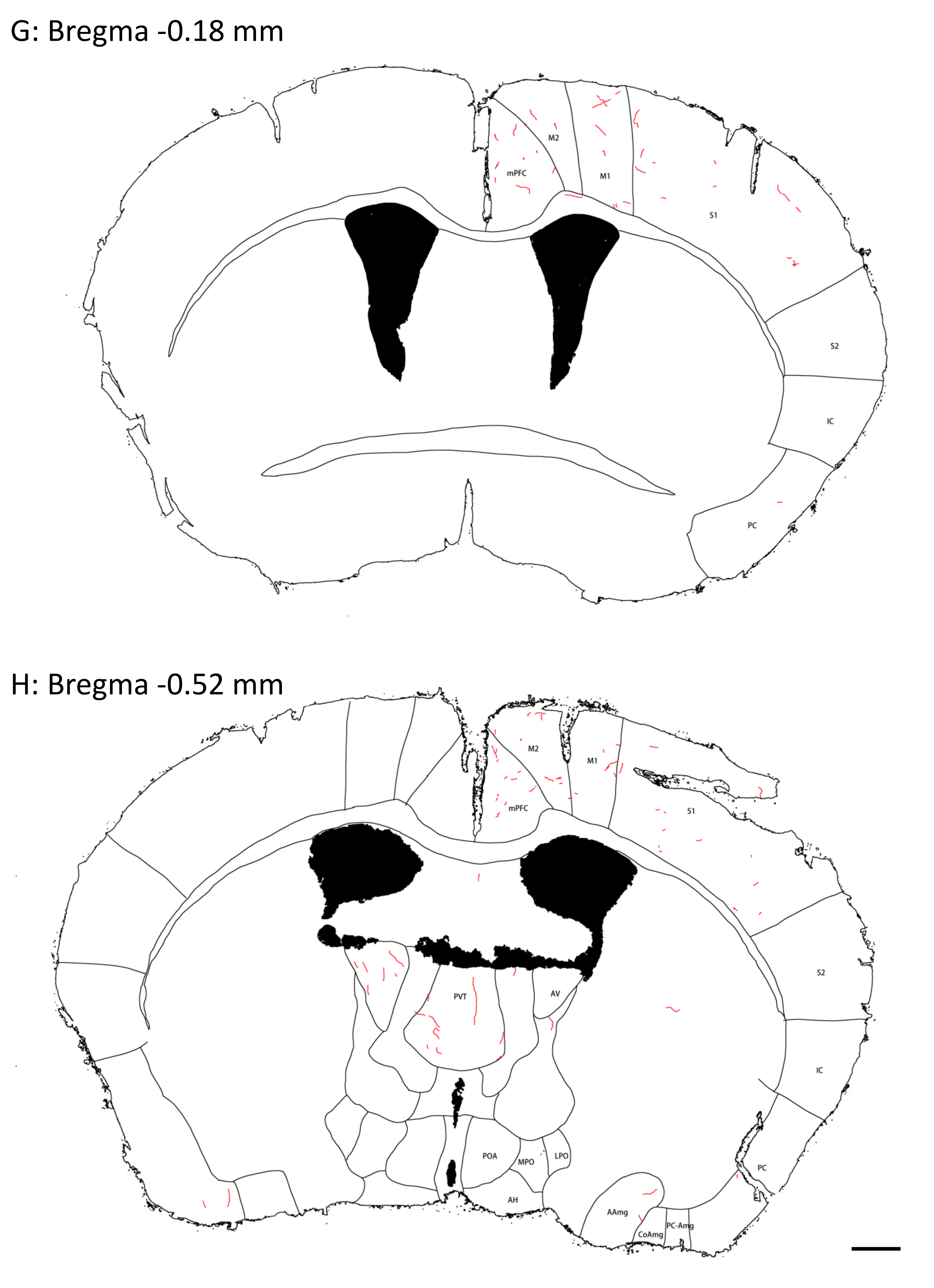
G: Bregma -0.18 mm
M2
M1
mPFC
S1
S2
IC
PC
H: Bregma -0.52 mm
M2
M1
mPFC
S1
PVT
AV
S2
IC
POA
MPO
LPO
PC
AH
AAmg
PC-Amg
CoAmg

I: Bregma -0.70 mm

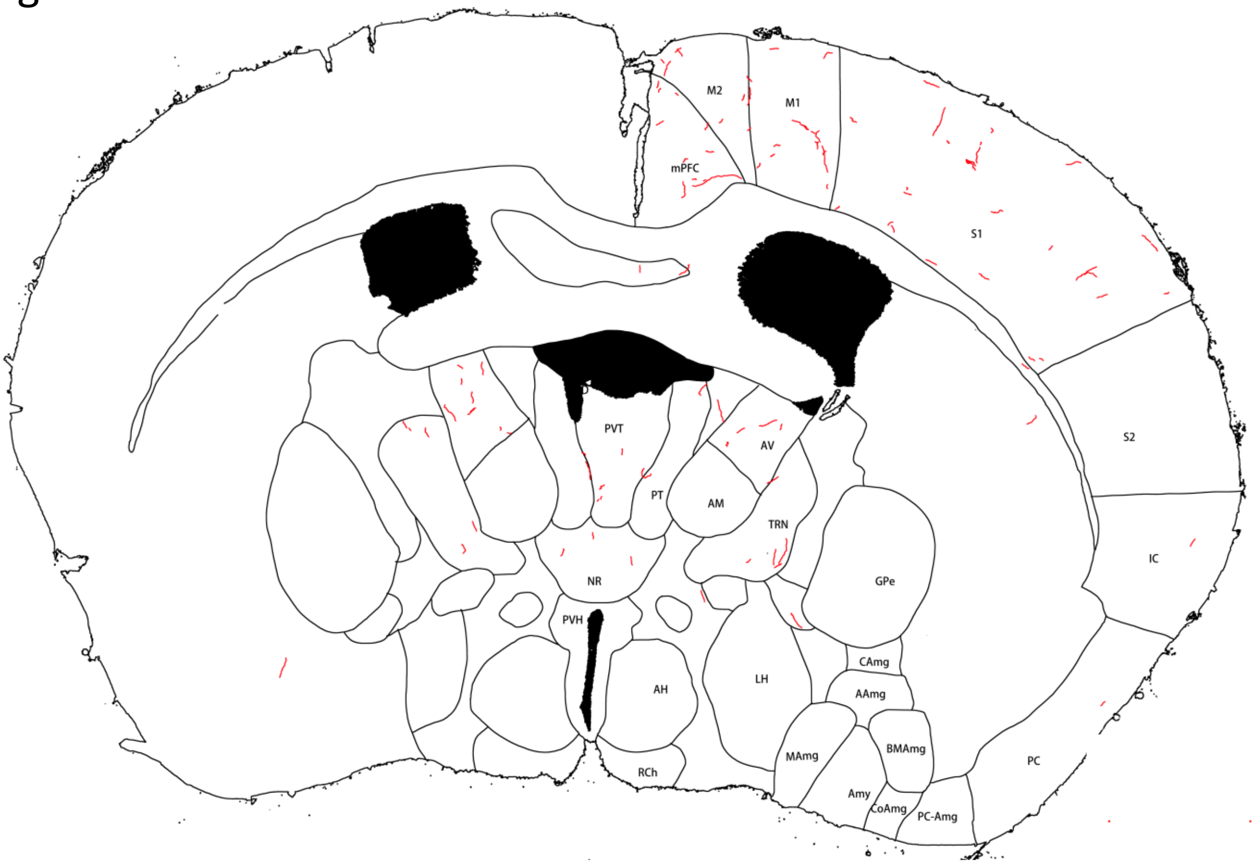


J: Bregma -1.14 mm

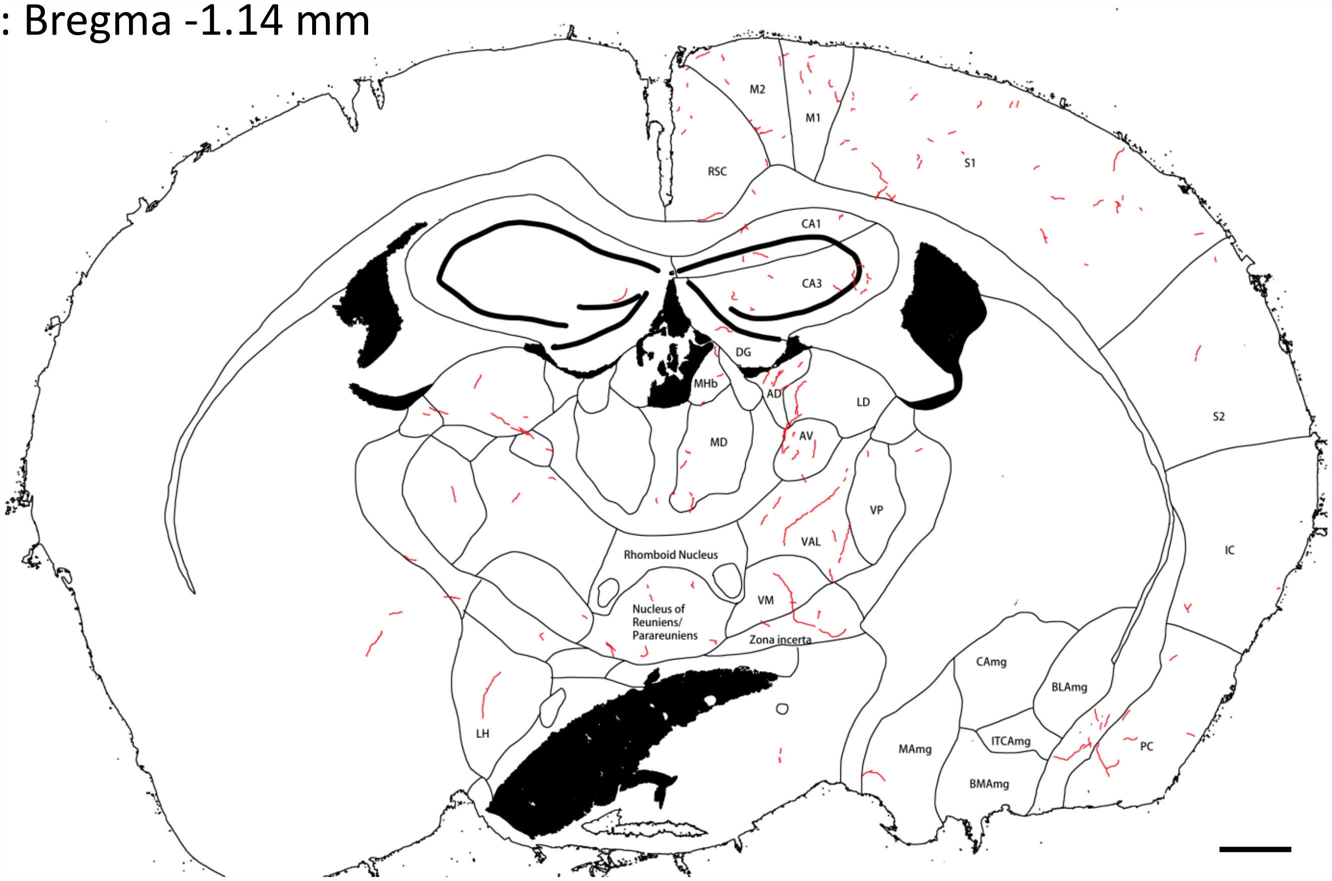

K: Bregma -1.53 mm

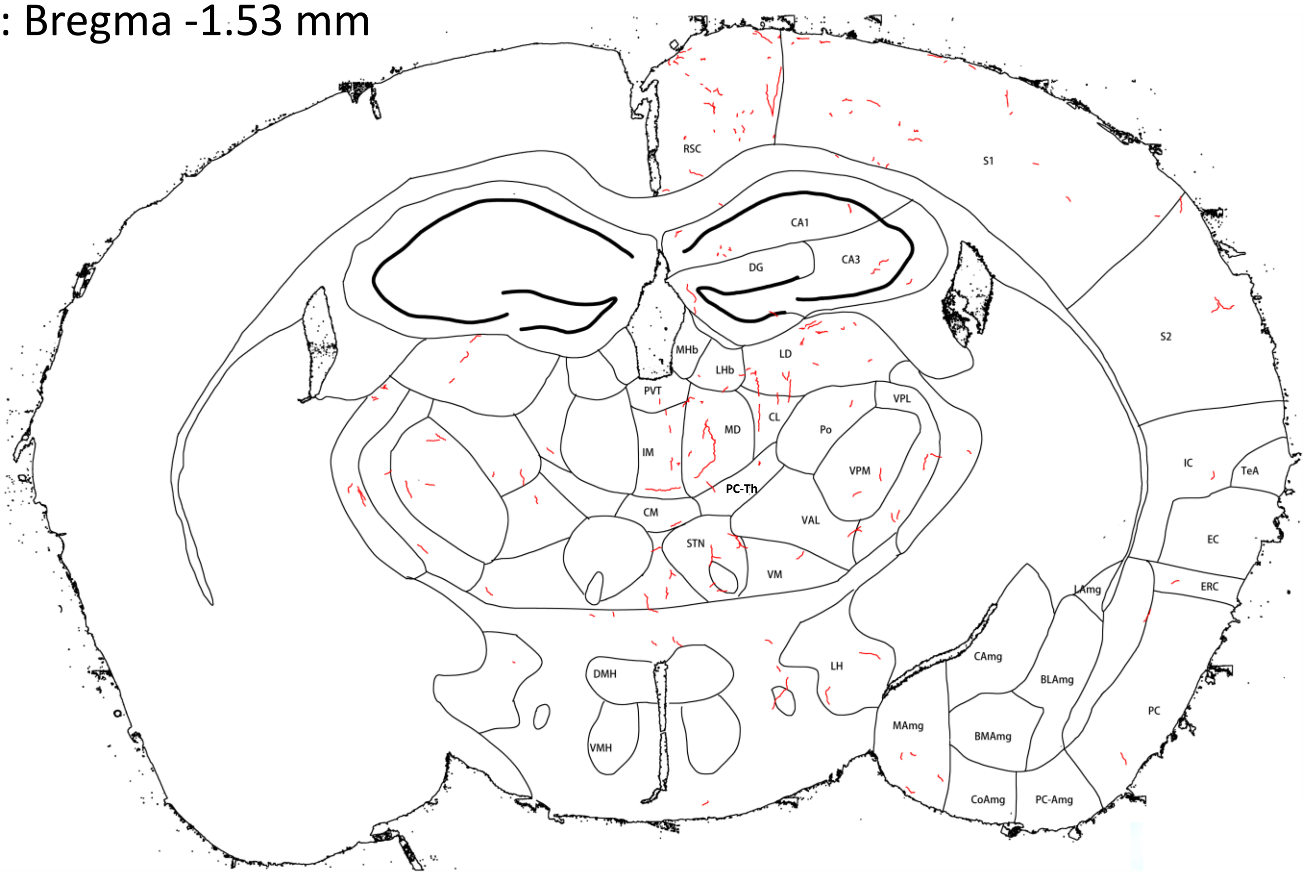


L: Bregma -1.85 mm

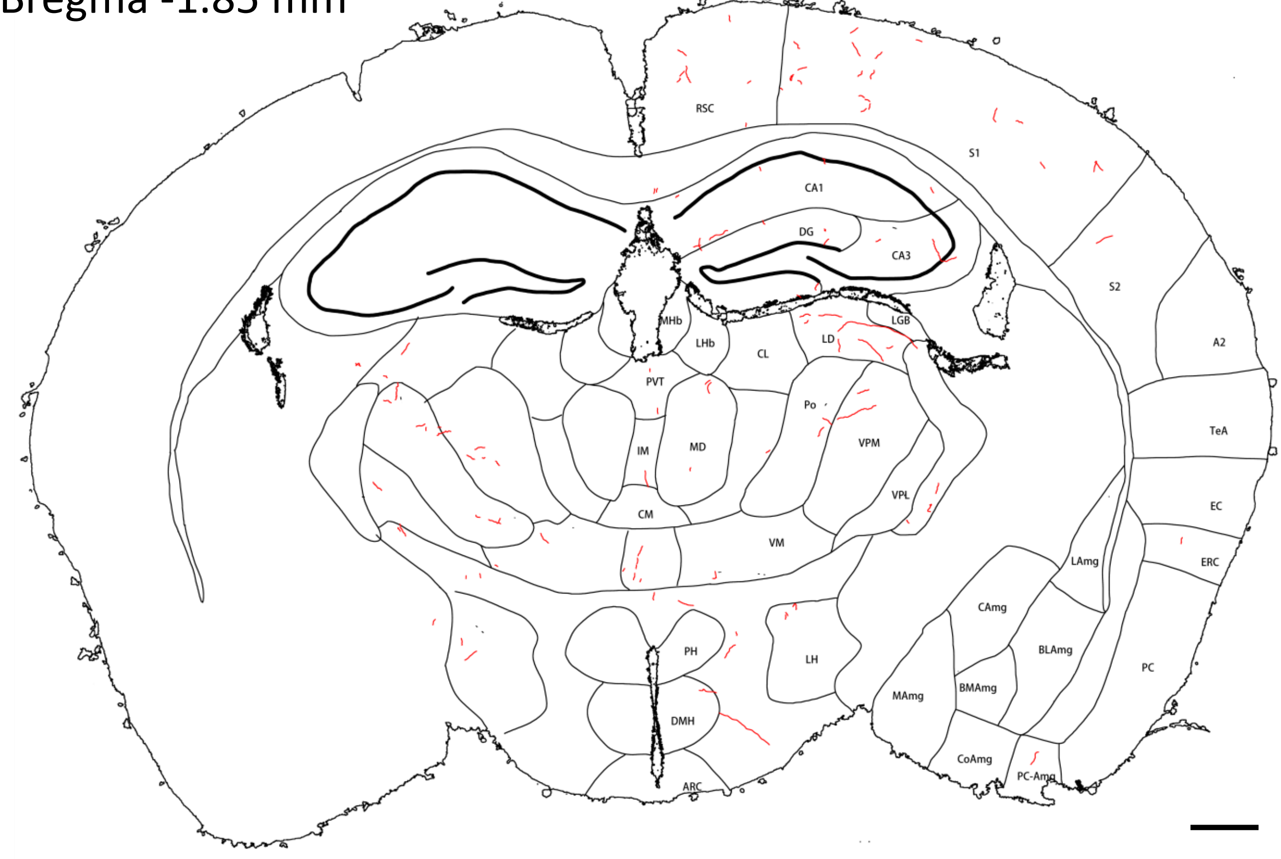

M: Bregma -2.19 mm

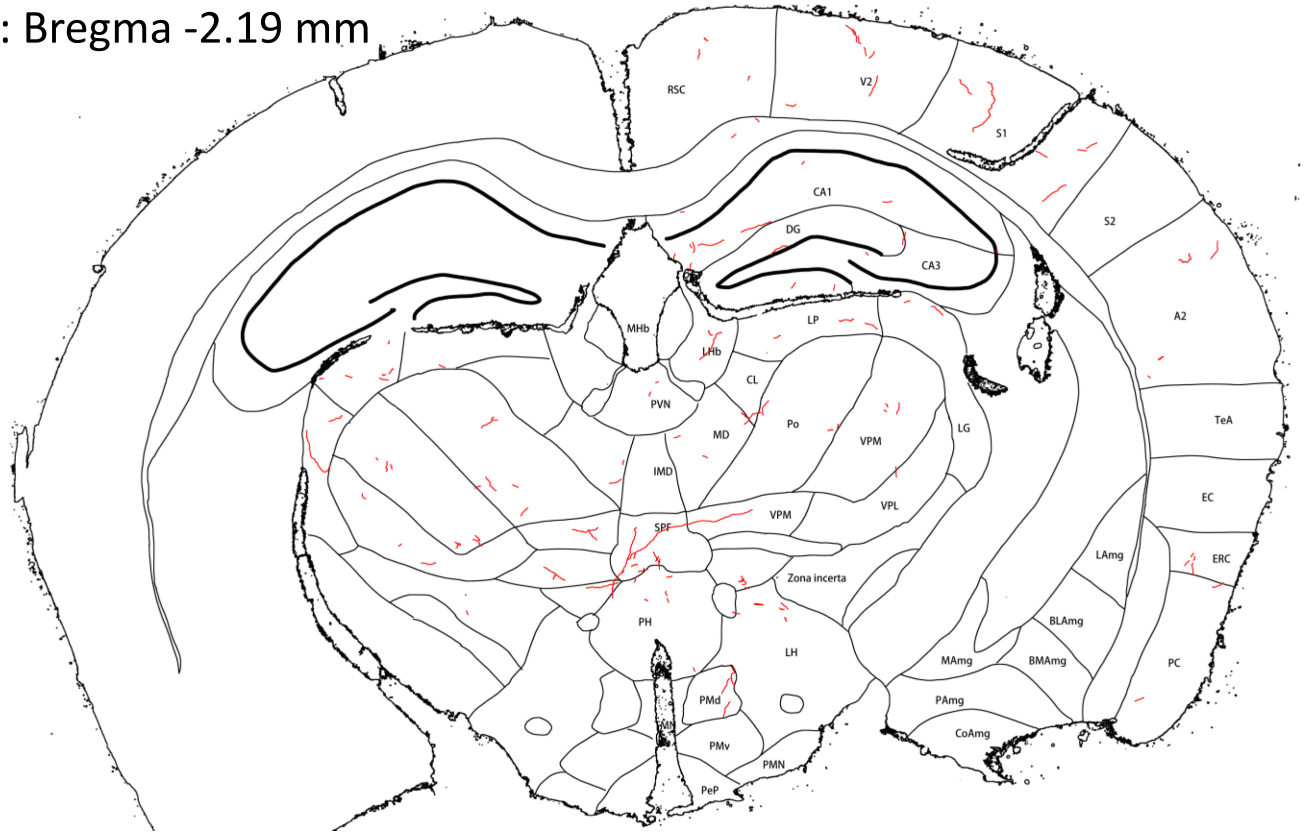


N: Bregma -2.49 mm

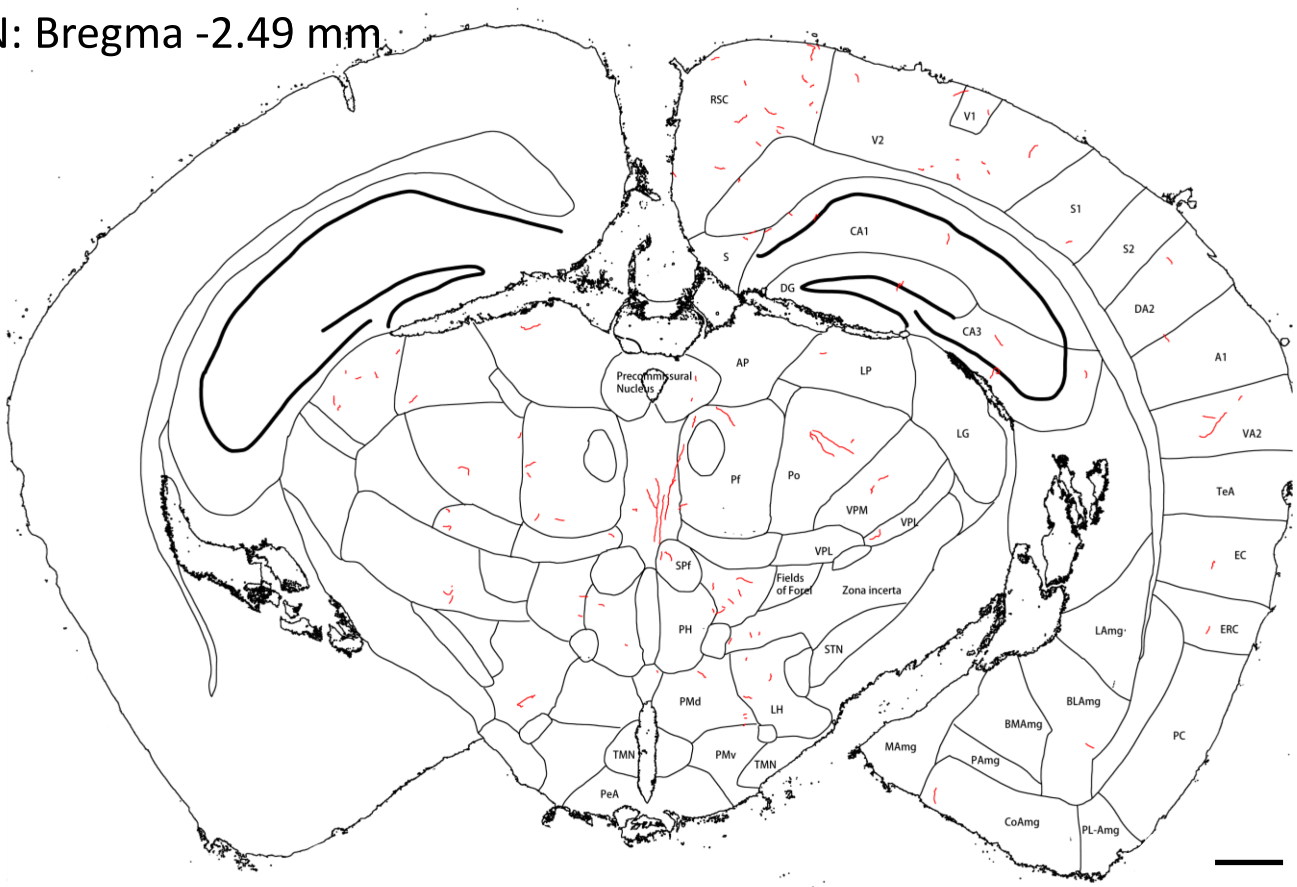

O: Bregma -2.70 mm

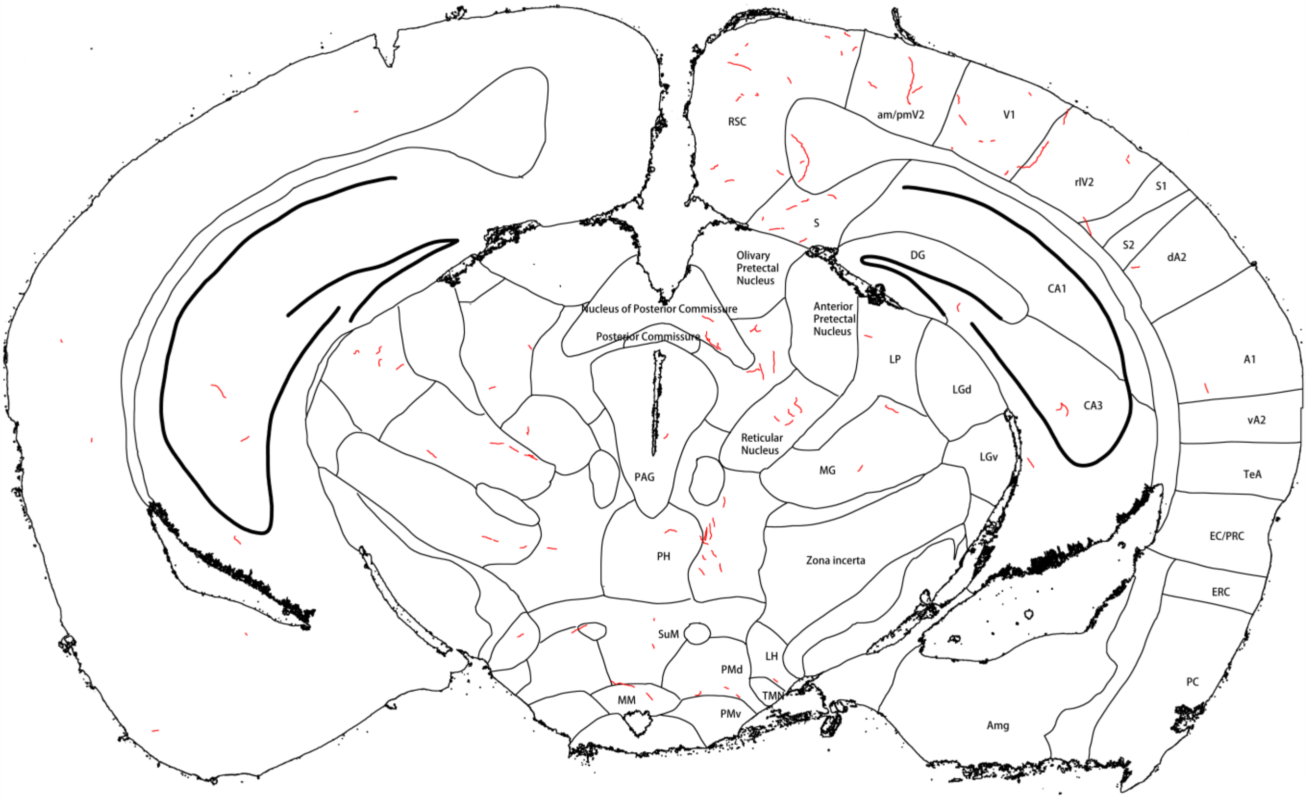


P: Bregma -3.02 mm

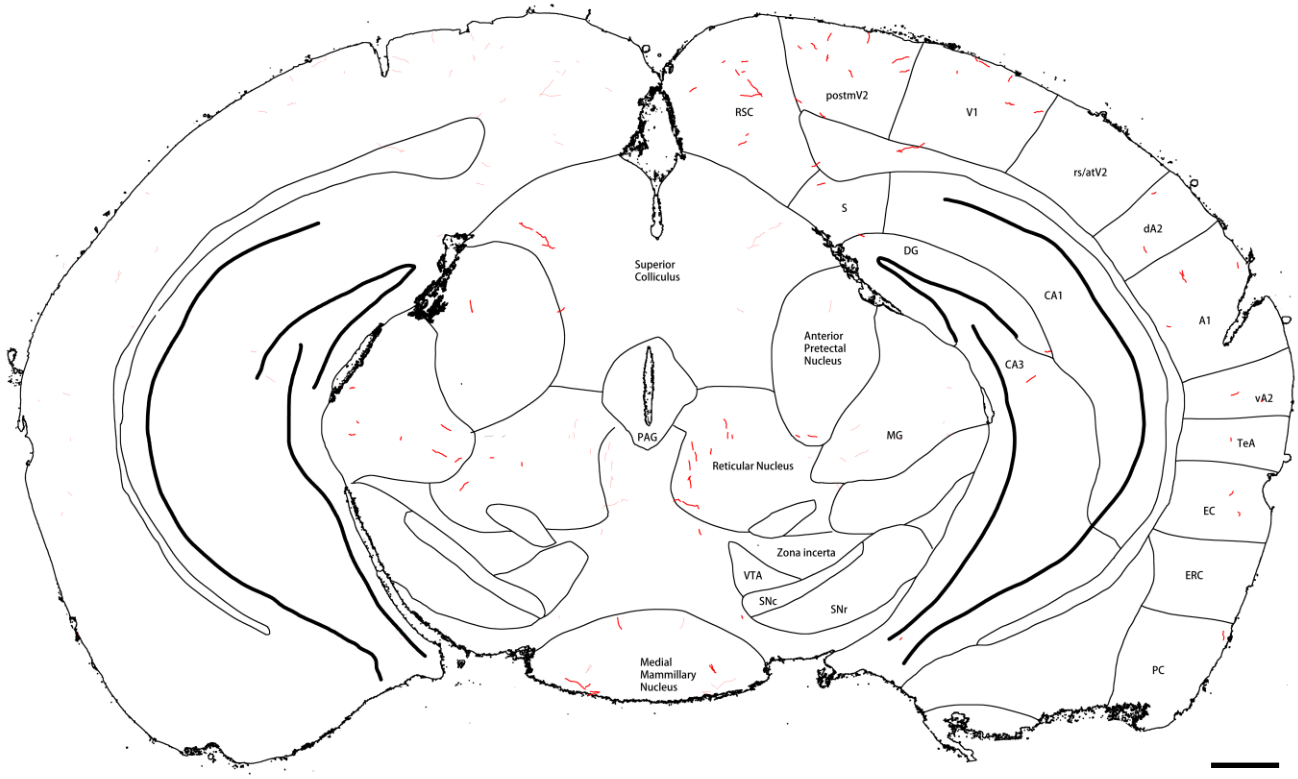

Q: Bregma -3.33 mm

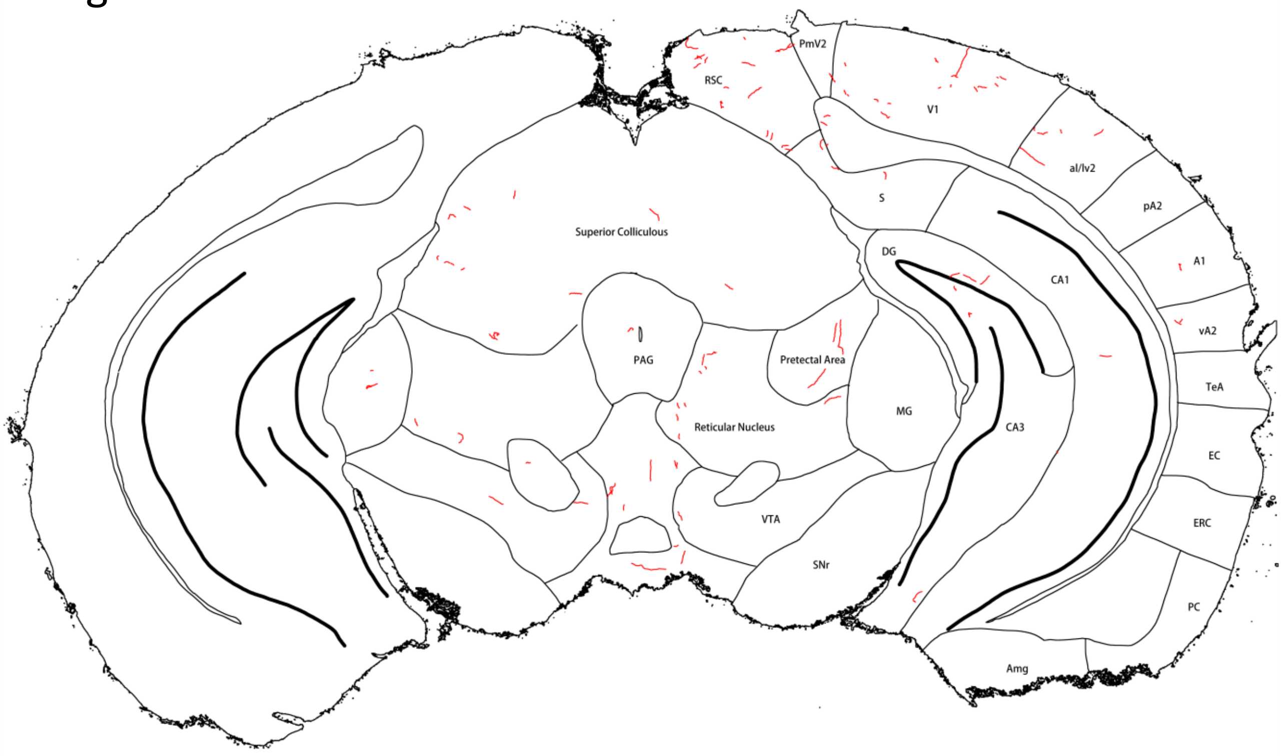


R: Bregma -3.71 mm

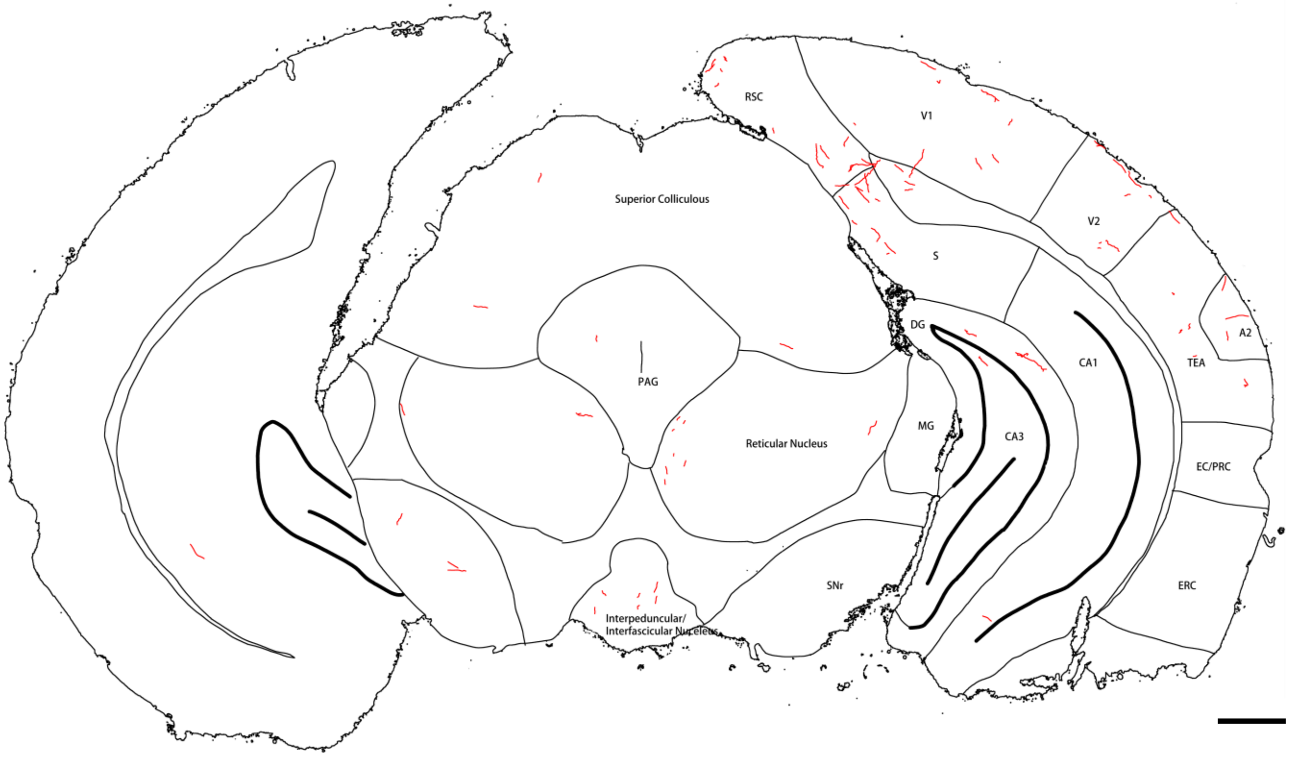


**Extended Figure 3C-1**. Tracing of NE fibers (red) in forebrain sections of mouse **#543**. Panels **A-F** are sections stained with FIHC; the others are sections stained with IHC/ABC. Yellow shadows in panels **B-D** mark the viral injection site with the core shown in panel **B**. Note that only sections where NE fibers are found are shown. Scale bar = 0.5 mm.

**List of brain region abbreviations:**

**A1**: Primary auditory cortex
**A2**: Secondary auditory cortex
**AAmg**: Anterior amygdalar nucleus
**ADHpc**: Anterodorsal hippocampus
**AH**: Anterior hypothalamus
**AM**: Anteromedial thalamic nucleus
**ARC**: Arcuate nucleus of the hypothalamus
**AV**: Anteroventral thalamic nucleus
**BLAmg**: Basolateral amygdalar nucleus
**BMAmg**: Basomedial amygdalar nucleus
**CA1**: Cornu ammonis area 1 (of the hippocampus)
**CA3**: Cornu ammonis area 3 (of the hippocampus)
**CAmg**: Central amygdalar nucleus
**CgC**: Cingulate cortex;
**CL**: Central lateral thalamic nucleus
**CoAmg**: Cortical amygdalar nucleus
**DG**: Dentate gyrus (of the hippocampus)
**DMH**: Dorsomedial hypothalamus
**EC**: Ectorhinal cortex
**ERC**: Entorhinal cortex
**IC**: Insular cortex
**IMD**: Intermedialdorsal thalamic nucleus
**ITCAmg**: Intercalated amygdalar nucleus
**LAmg**: Lateral amygdalar nucleus
**LD**: Lateral dorsal thalamic nucleus
**LGN**: Lateral geniculate nucleus
**LH**: Lateral hypothalamus
**LHb**: Lateral habenula
**LPO**: Lateral preoptic area
**M1**: Primary motor cortex
**M2**: Secondary motor cortex
**MAmg**: Medial amygdalar nucleusr
**MD**: Mediodorsal thalamic nucleus
**MGN**: Medial geniculate nucleus
**MHb**: Medial habenula
**MPO**: Medial preoptic area
**NR**: Nucleus reuniens (of the thalamus)
**PAmg**: Posterior amygdalar nucleus
**PC**: Piliform cortex
**PC-Amg**: Pliform-amygdalar area
**PC-Th**: Paracentral thalamic nucleus
**PDHpc**: Posterodorsal hippocampus
**PeA**: Periventricular hypothalamic nucleus
**Pf**: Parafascicular thalamic nucleus
**PH**: Posterior hypothalamus
**PMd**: Dorsal premammillary nucleus
**PMv**: Ventral premammillary nucleus
**Po**: Posterior thalamic nucleus
**POA**: Preoptic area
**PRC**: Perirhinal cortex
**PT**: Paratenial thalamus
**PVH**: Paraventricular hypothalamus
**PVT**: Paraventricular thalamus
**RCh**: Retrochiasmatic area
**RSC**: Retrosplenial cortex
**S**: Subiculum (part of the hippocampal formation)
**S1**: Primary somatosensory cortex
**S2**: Secondary somatosensory cortex
**SNc**: Substantia nigra, pars compacta
**SNr**: Substantia nigra, pars reticulata
**SPF**: Subparafascicular thalamic nucleus
**STN**: Subthalamic nucleus
**TeA**: Temporal association area
**TMN**: Tuberomammillary nucleus
**TRN**: Thalamic reticular nucleus
**V1**: Primary visual cortex
**V2**: Secondary visual cortex
**VAL**: Ventral anterior-lateral thalamic complex
**VHpc**: Ventral hippocampus
**VM**: Ventromedial thalamic nucleus
**VPL**: Ventral posterolateral thalamic nucleus
**VPM**: Ventral posteromedial thalamic nucleus
**VTA**: Ventral tegmental area
**ZI**: Zona incerta
