## Supplementary Figure 2 for "Axon Collateral Pattern of a Sparse Locus Coeruleus Norepinephrine Neuron in Mouse Cerebral Cortex"

NE fibers (red) in forebrain sections of **Mouse #544**.

A: Bregma 1.30 mm

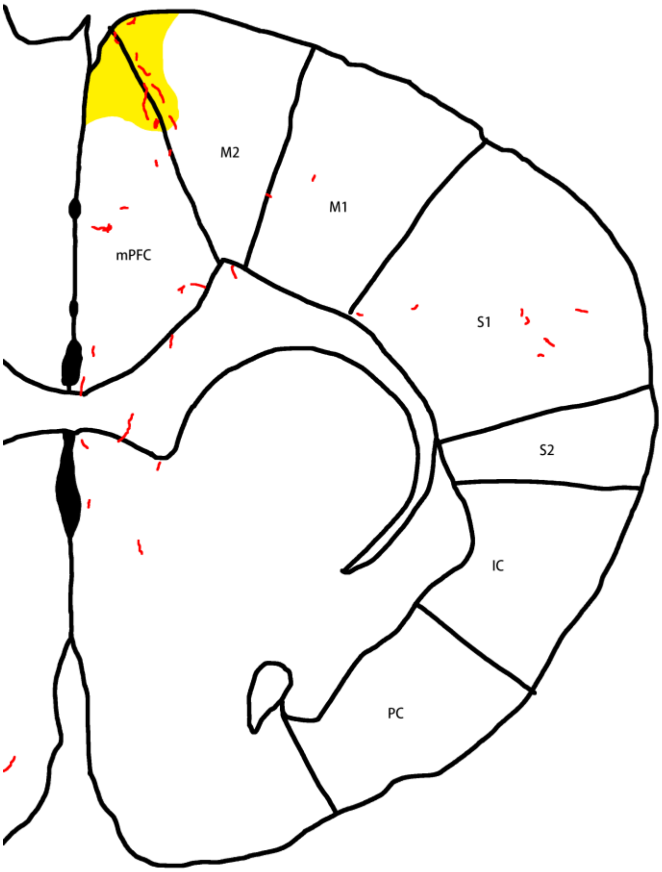


B: Bregma 0.98 mm
(Core injection site)

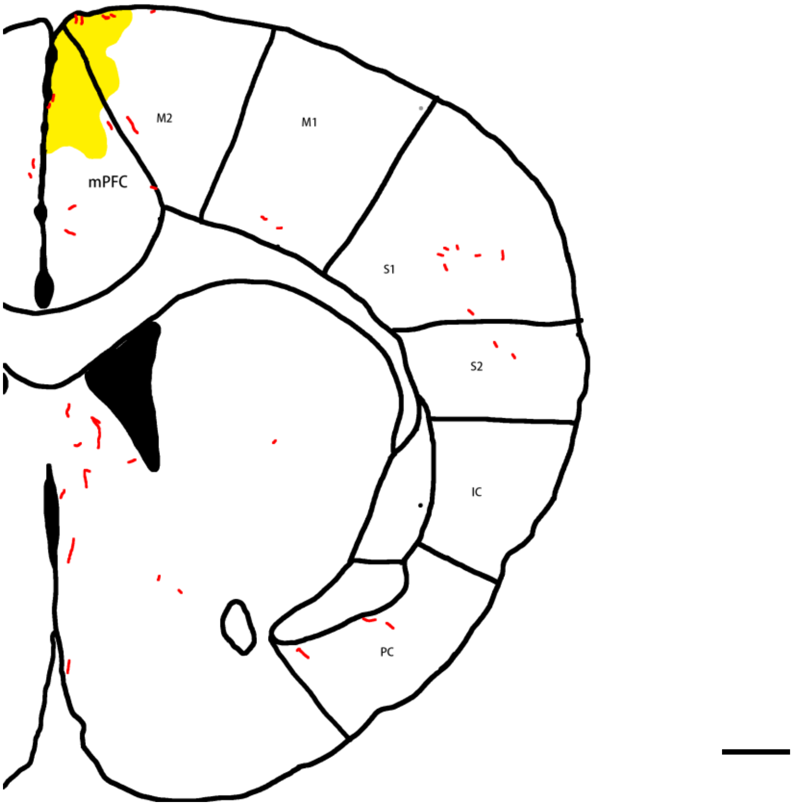

C: Bregma 0.78 mm

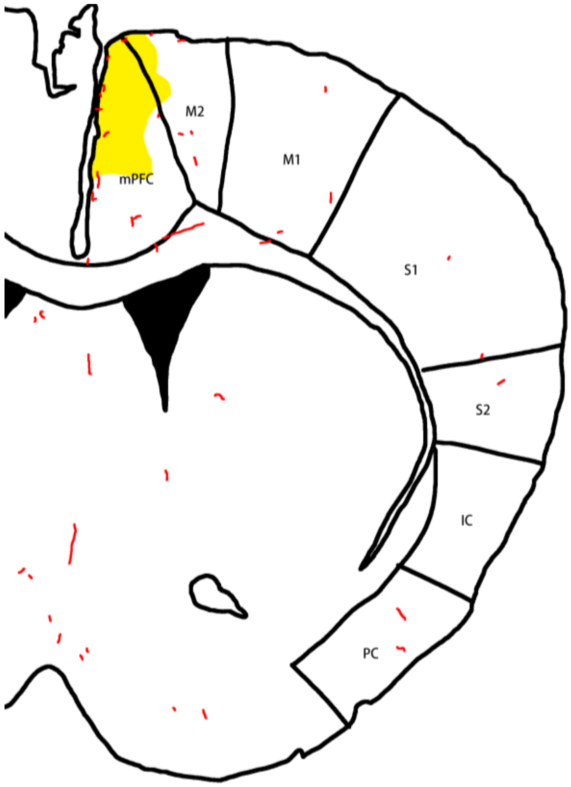


D: Bregma 0.63 mm

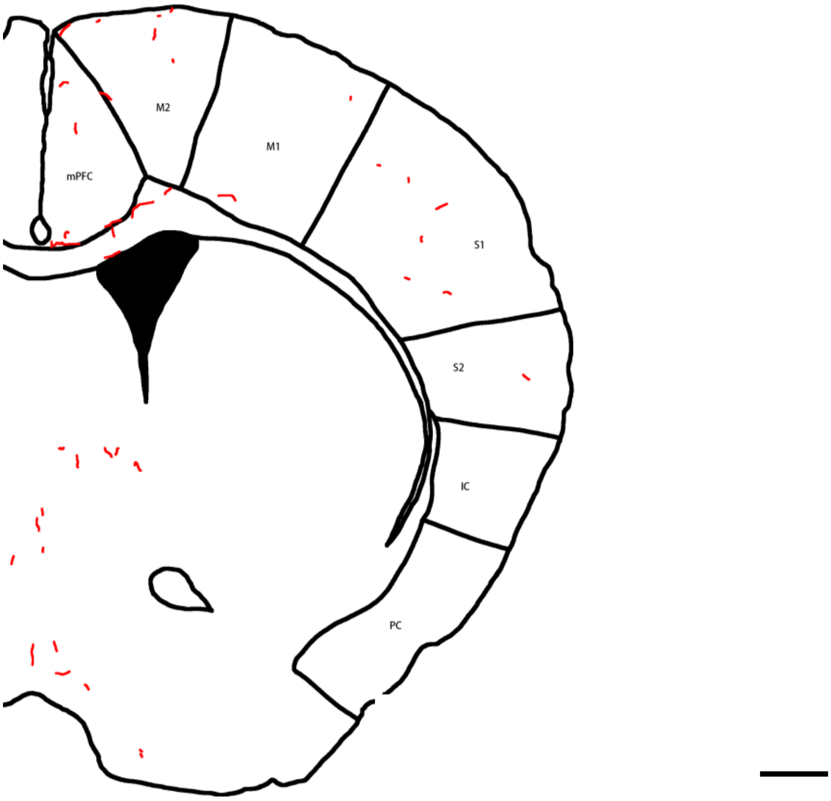

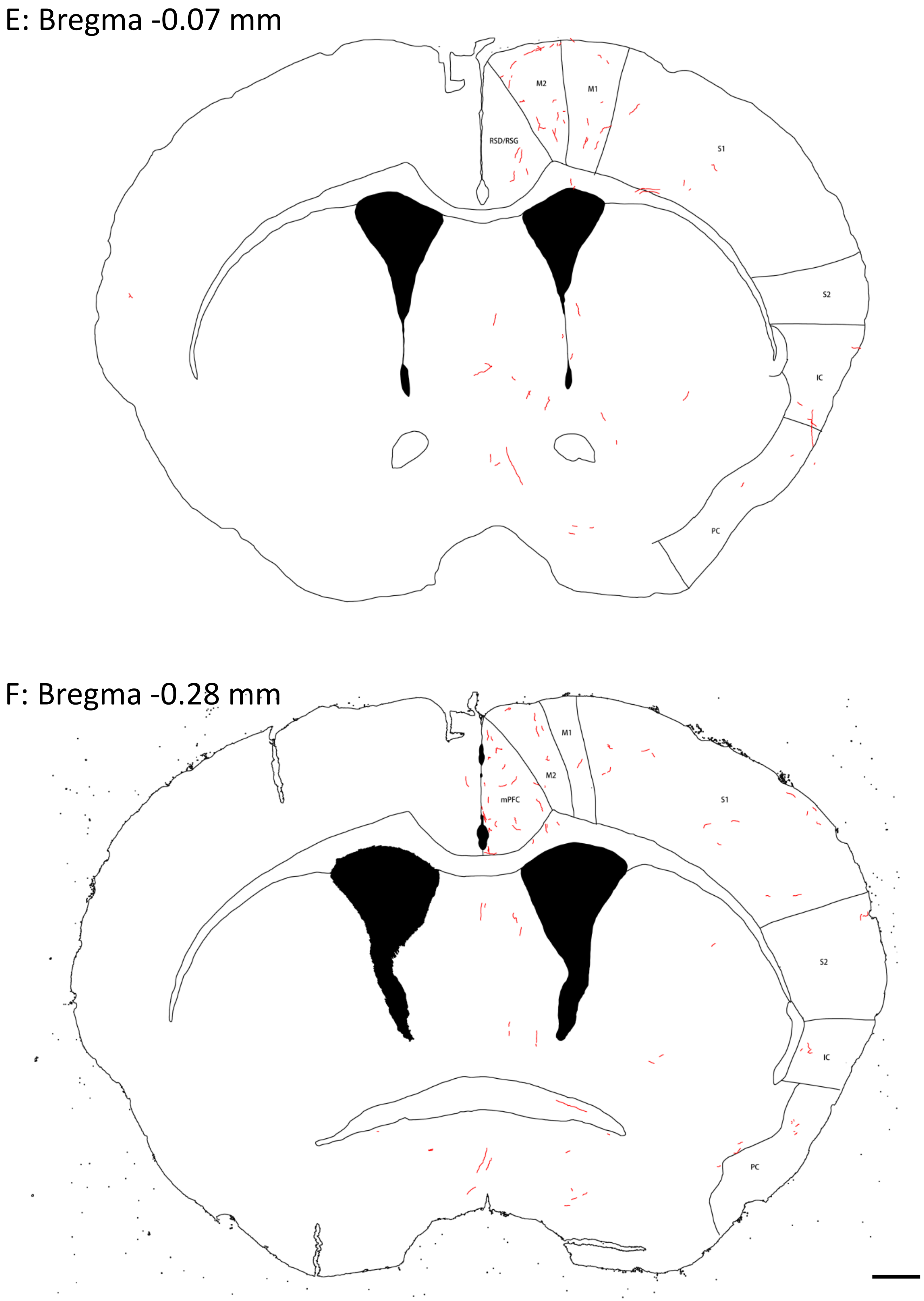
E: Bregma -0.07 mm
M2
M1
RSD/RSG
S1
S2
IC
PC
F: Bregma -0.28 mm
M1
M2
mPFC
S1
S2
IC
PC

G: Bregma -0.60 mm

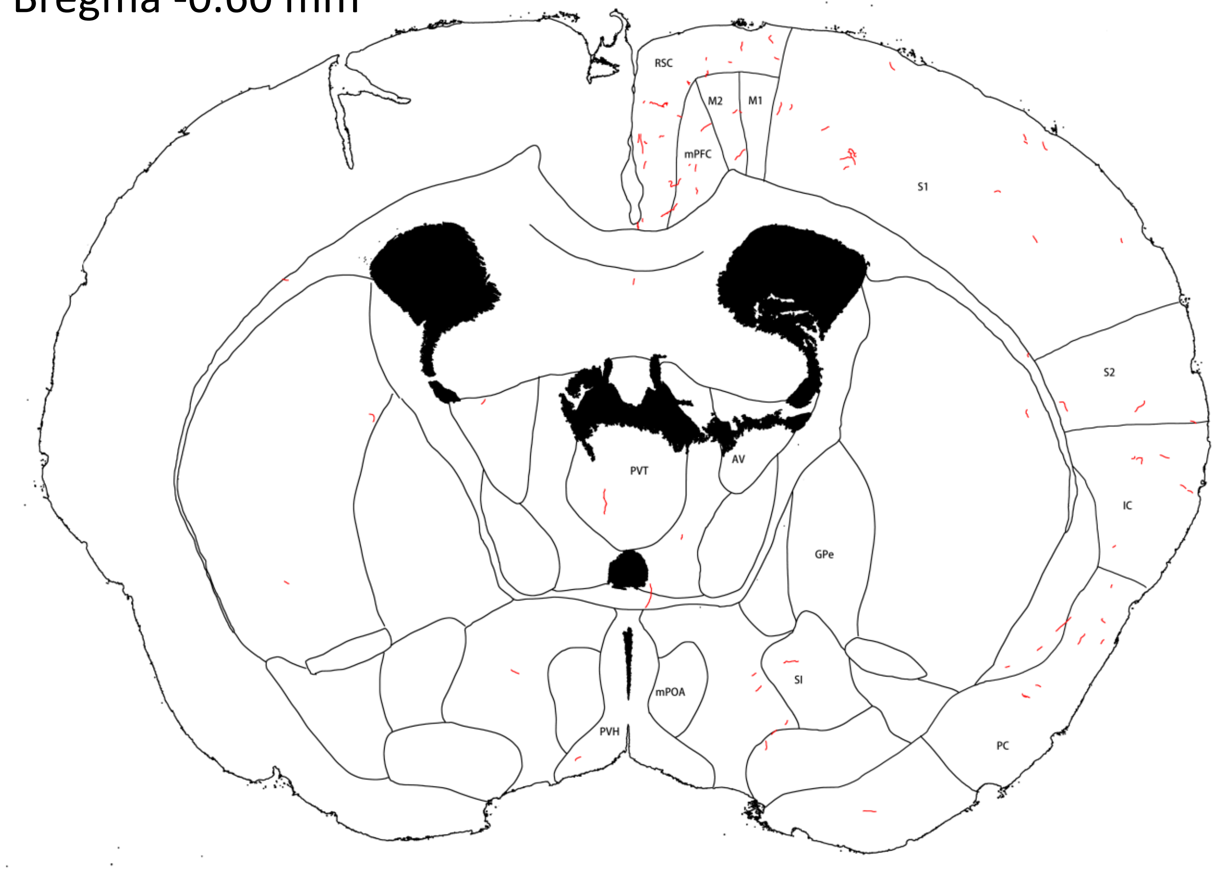


H: Bregma -0.79 mm

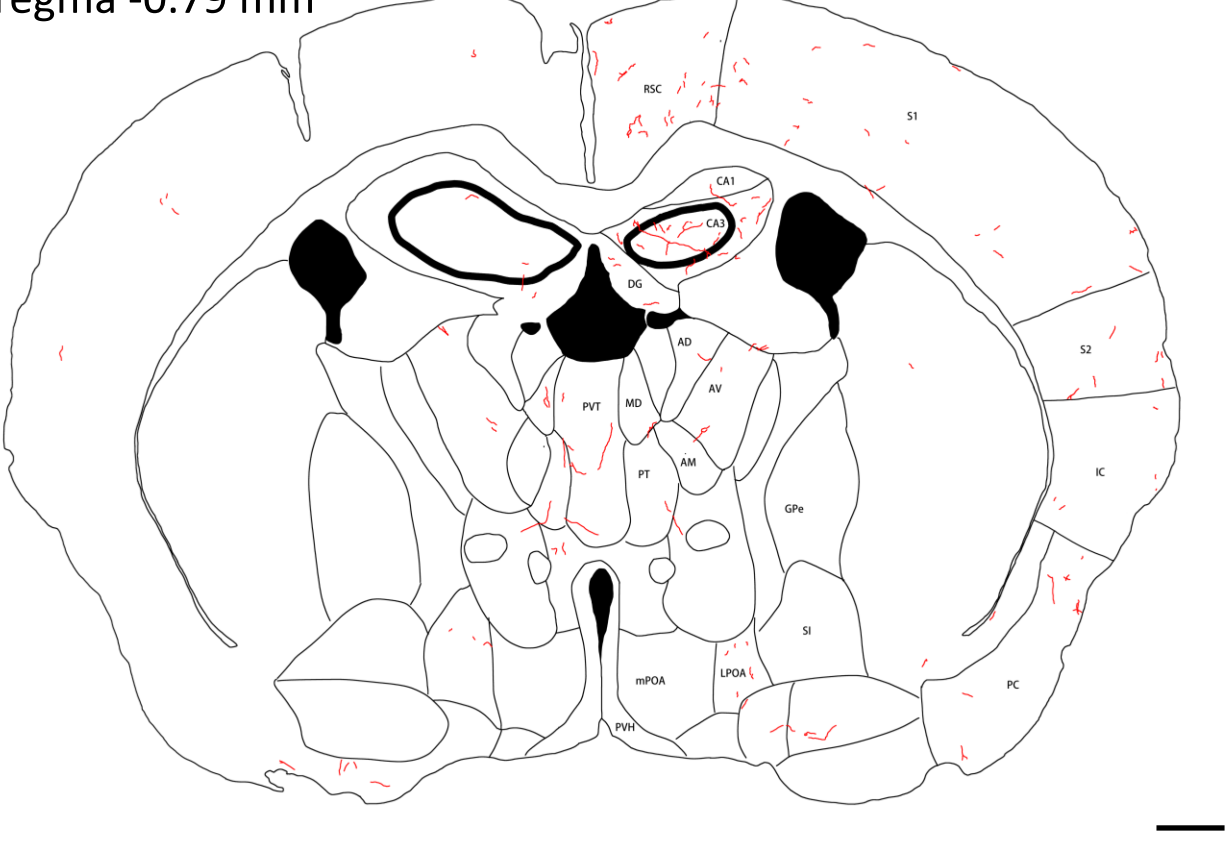

I: Bregma -1.08 mm

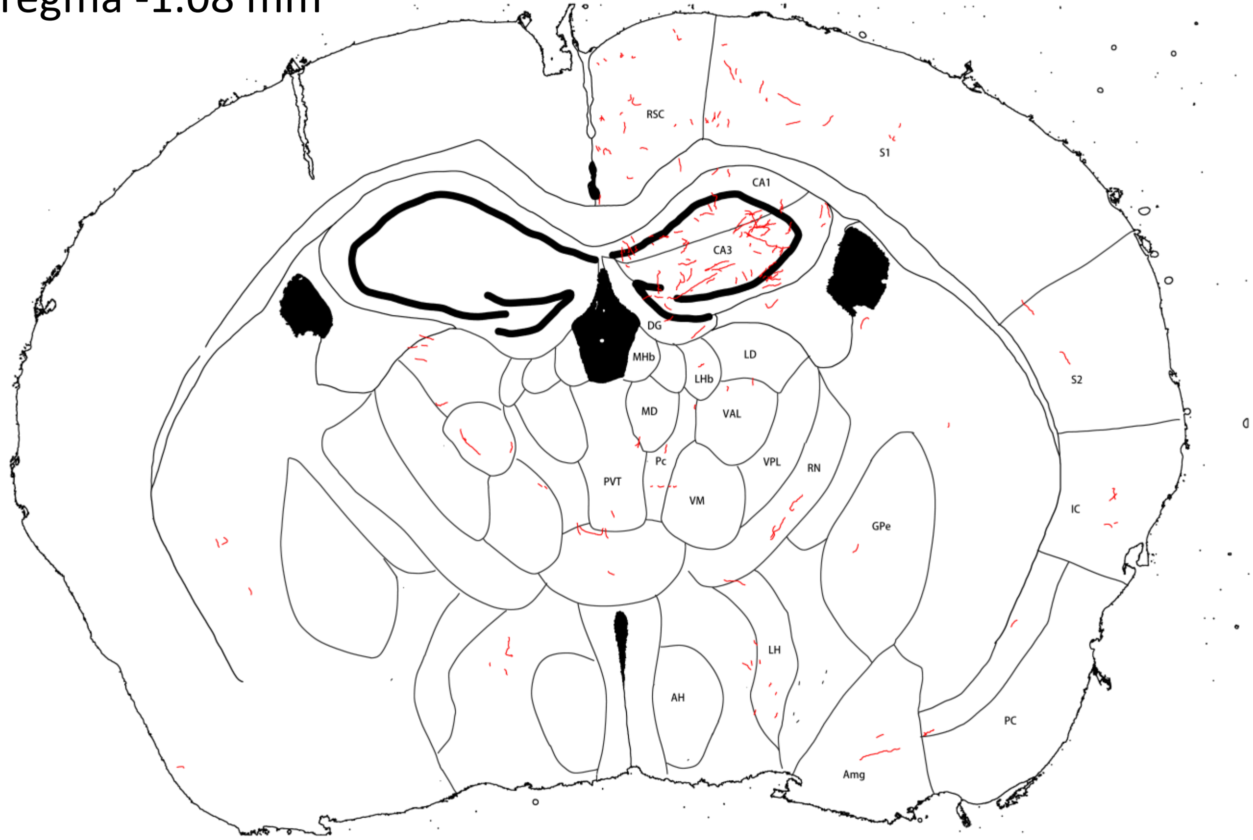


J: Bregma -1.48 mm

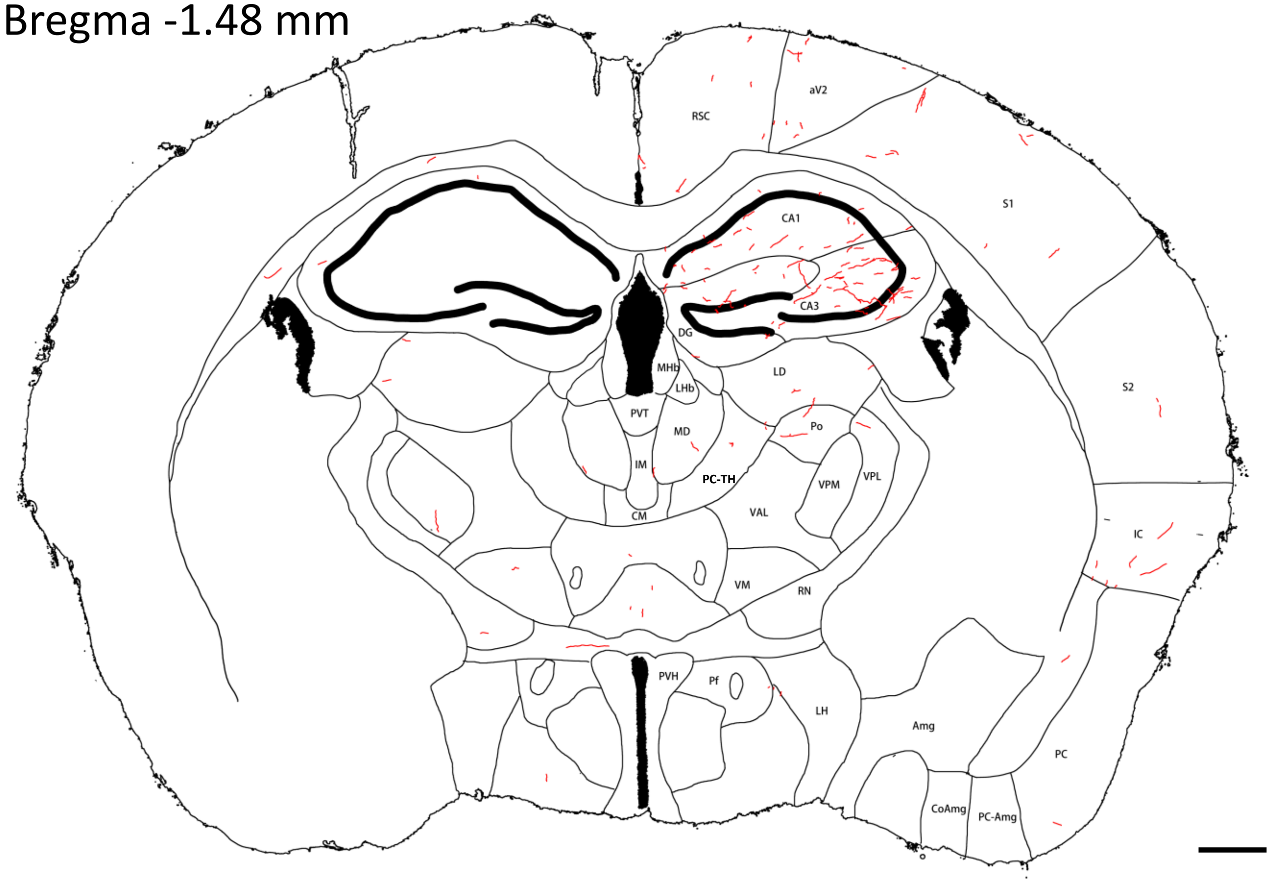

K: Bregma -1.92 mm

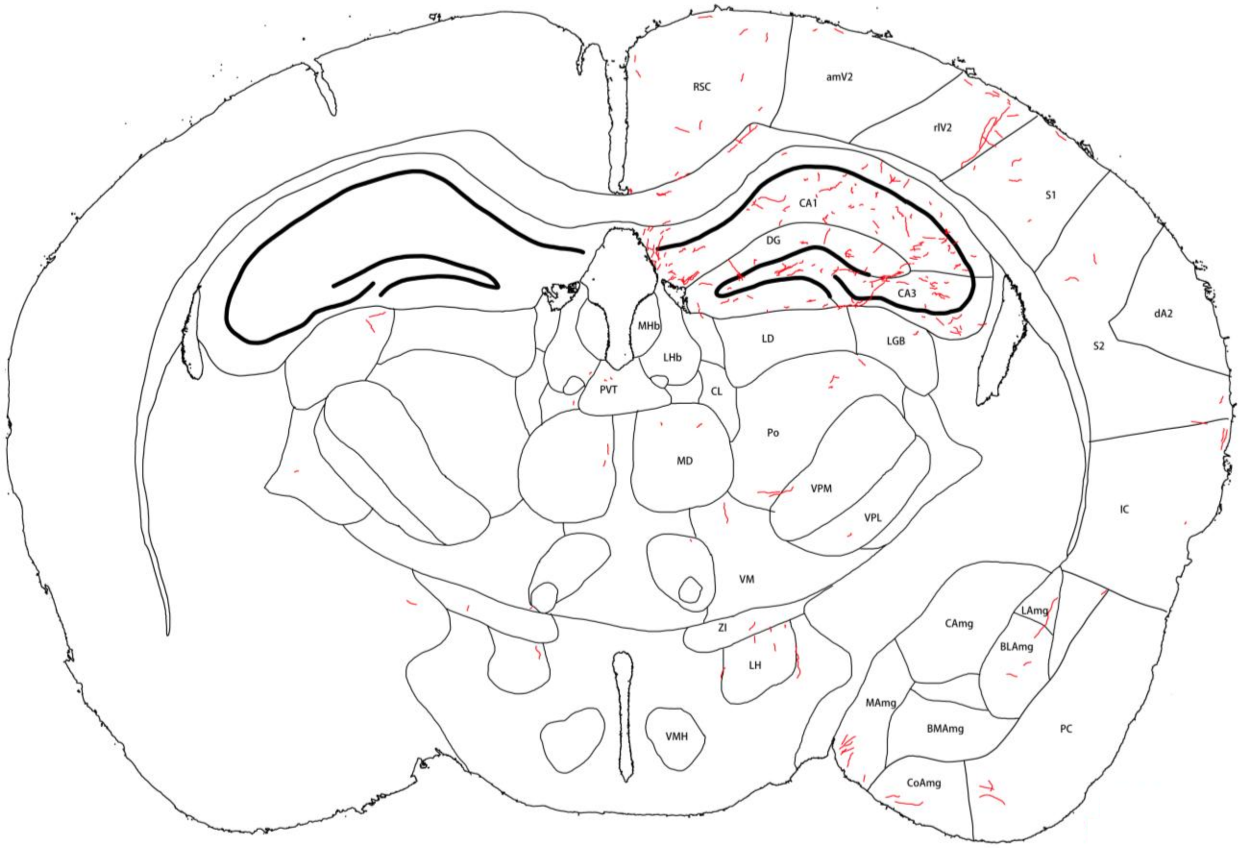


L: Bregma -2.25 mm

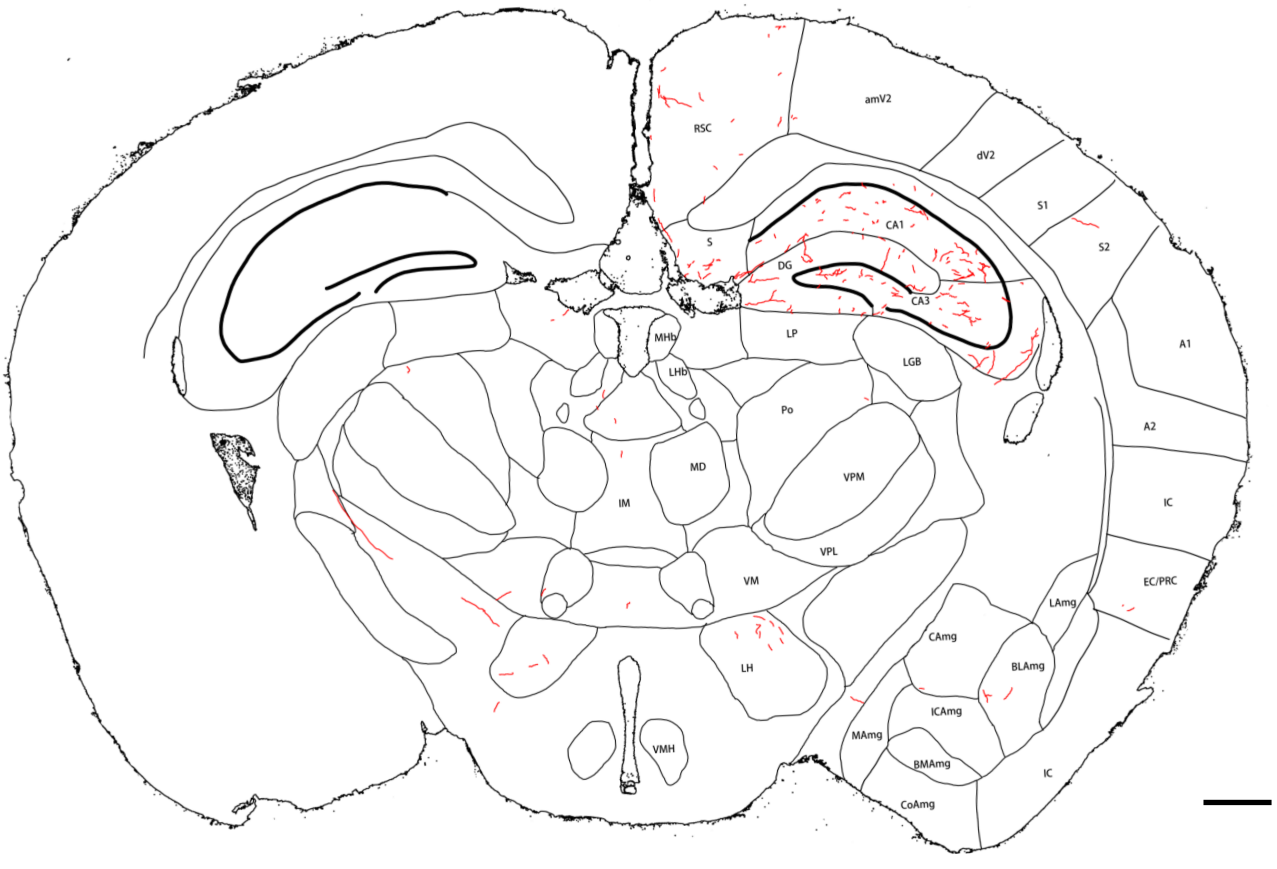

M: Bregma -2.45 mm

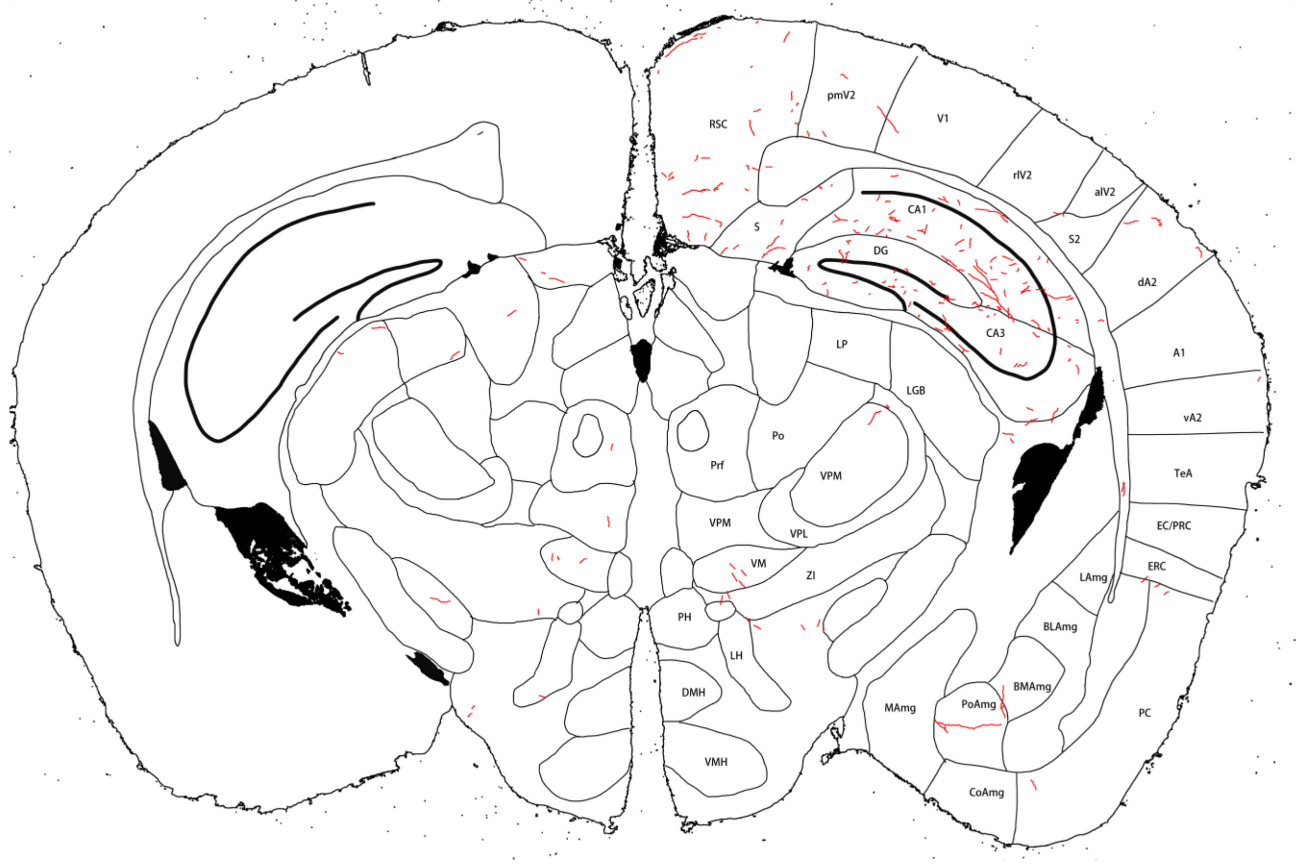


N: Bregma -2.75 mm

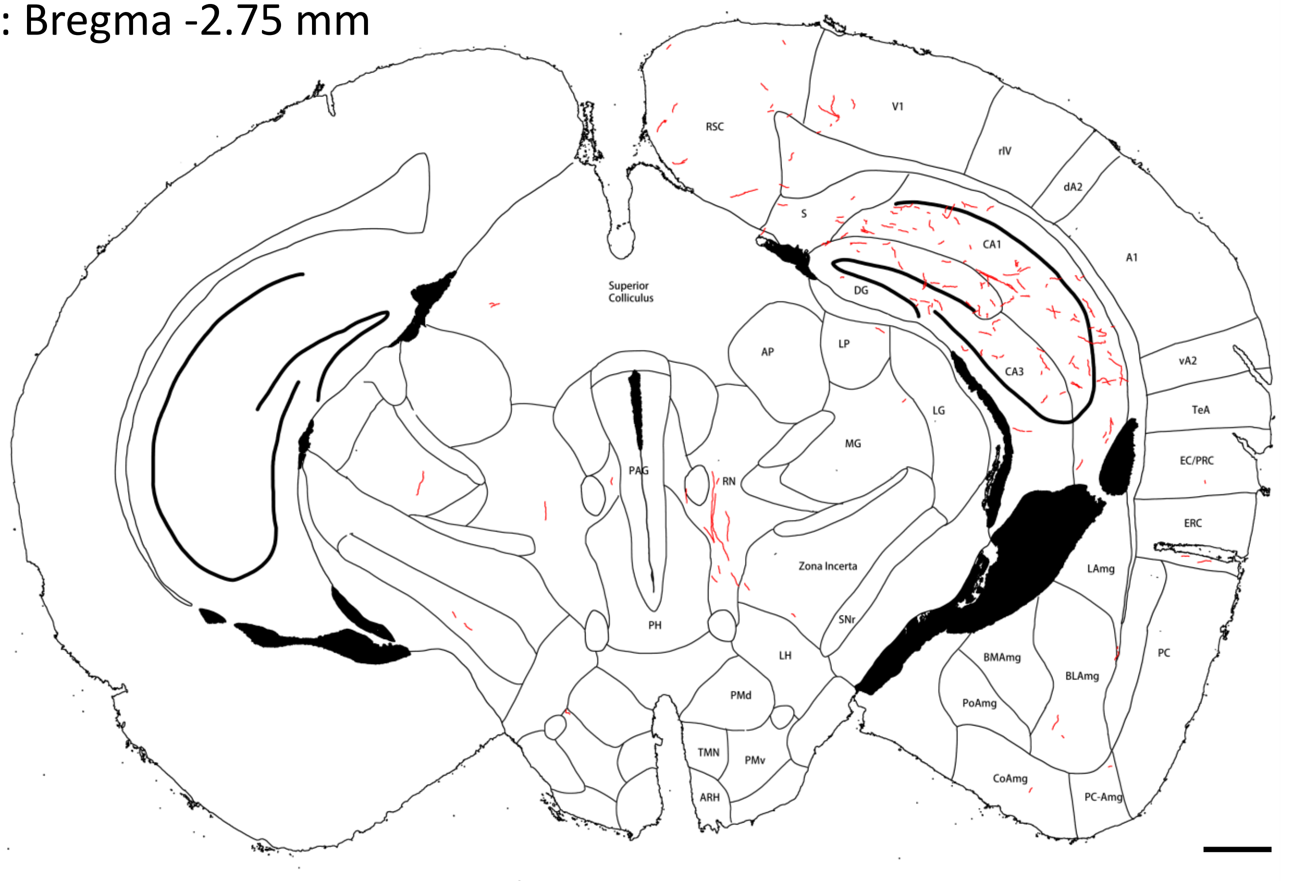

O: Bregma -3.07 mm

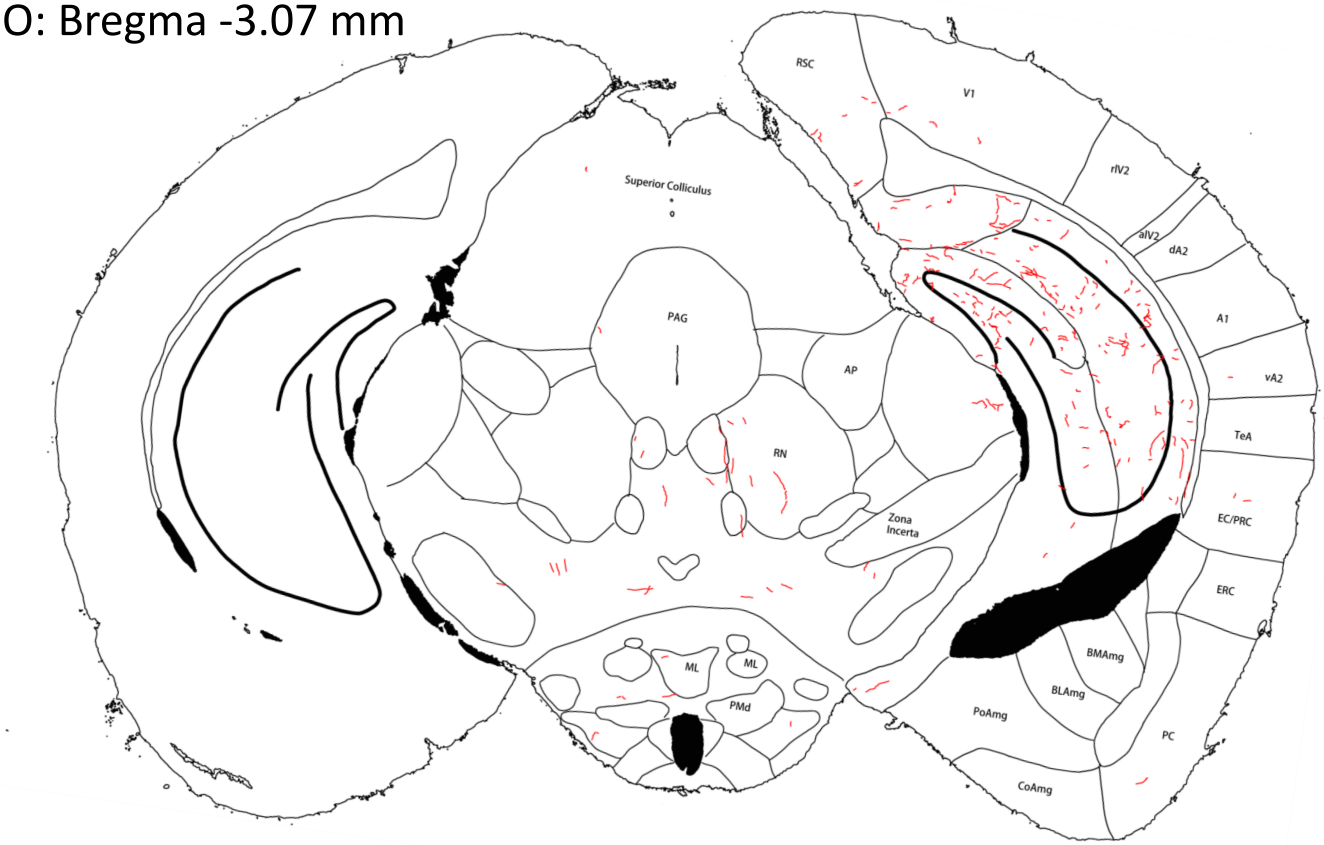


P: Bregma -3.29 mm

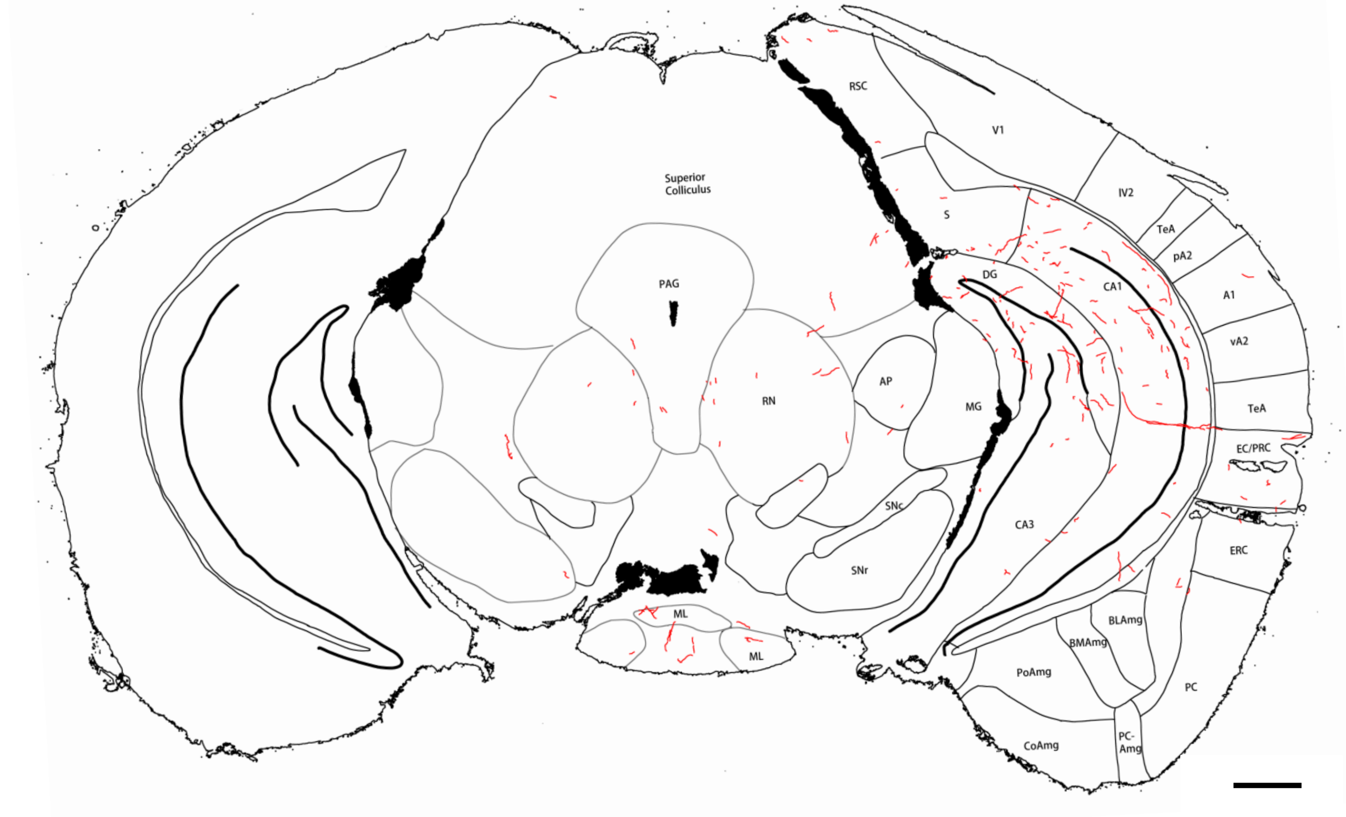

Q: Bregma -3.61 mm

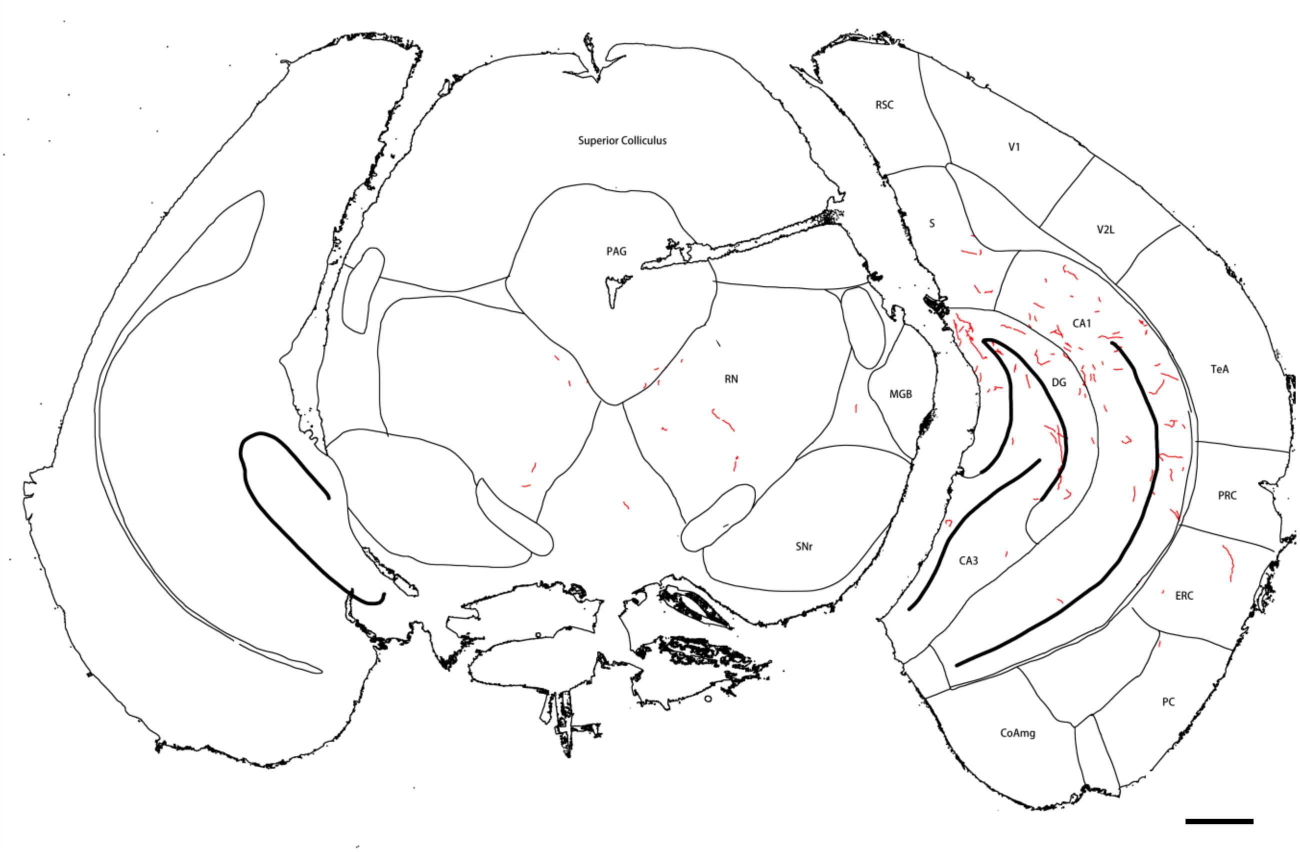


**Extended Figure 3C-2**. Tracing of NE fibers (red) in forebrain sections of mouse **#544**. Panels **A-D** are sections stained with FIHC; the others are sections stained with IHC/ABC. Yellow shadows in panels **A-C** mark the viral injection site with the core shown in panel **B**. Note that only sections where NE fibers are found are shown. Scale bar = 0.5 mm.
