## Supplementary Figure 3 for "Axon Collateral Pattern of a Sparse Locus Coeruleus Norepinephrine Neuron in Mouse Cerebral Cortex"

NE fibers (red) in forebrain sections of **Mouse #513**.

A: Bregma 0.51 mm

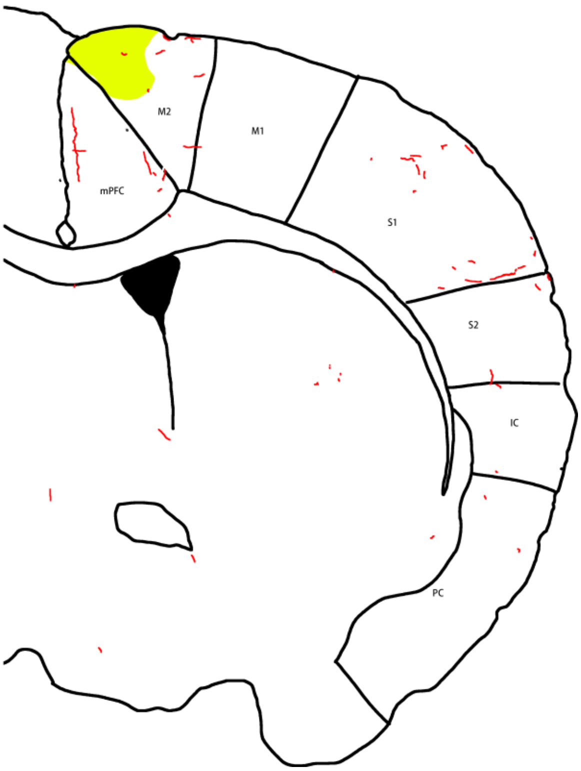


B: Bregma 0.26 mm
(Core injection site)

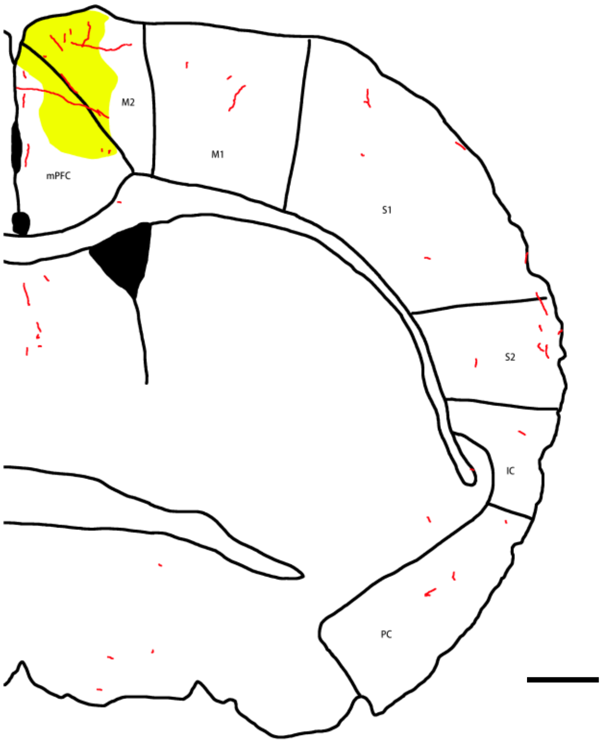

C: Bregma -0.13 mm

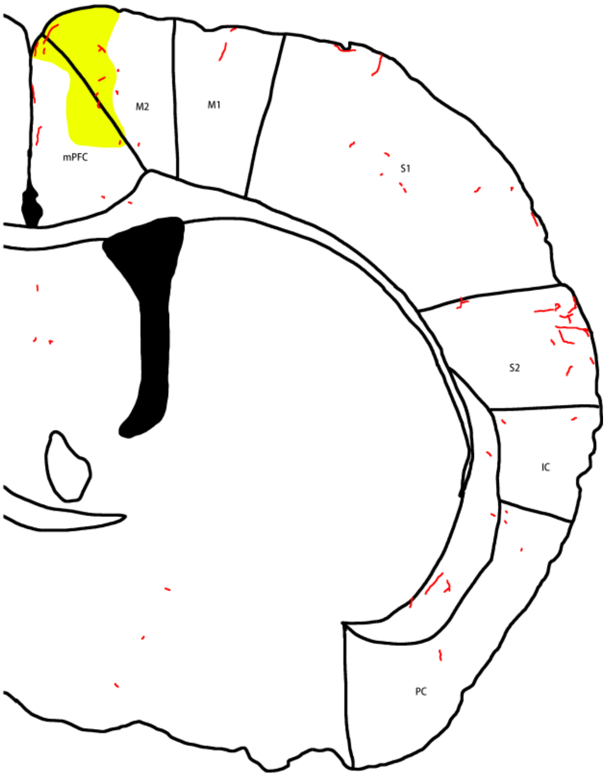


D: Bregma -0.16 mm

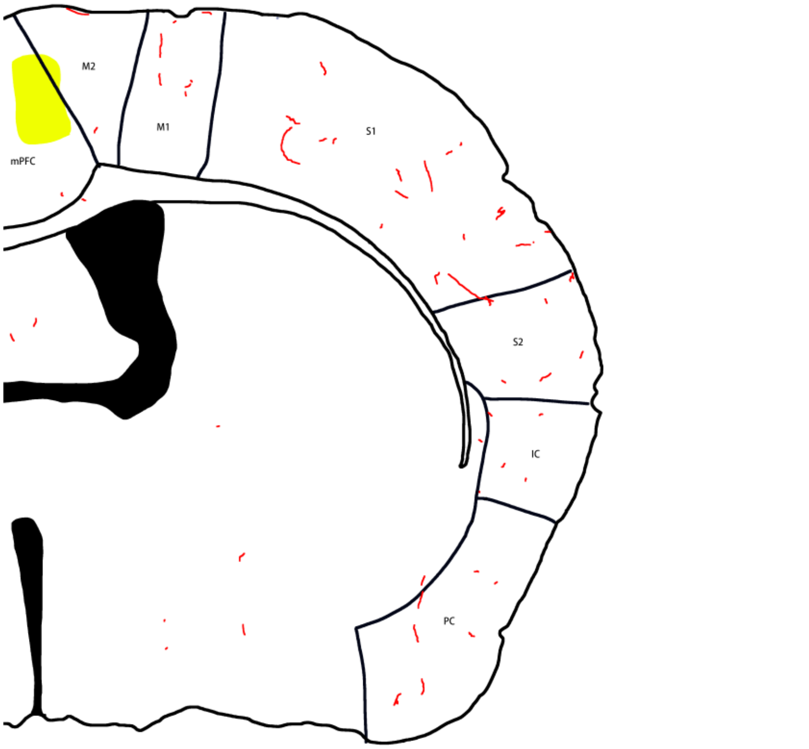

E: Bregma -0.51 mm

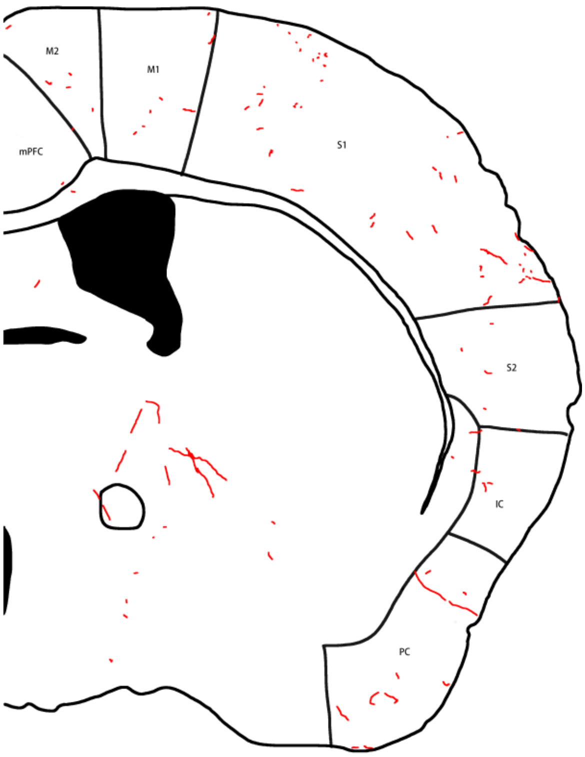


F: Bregma -0.68 mm

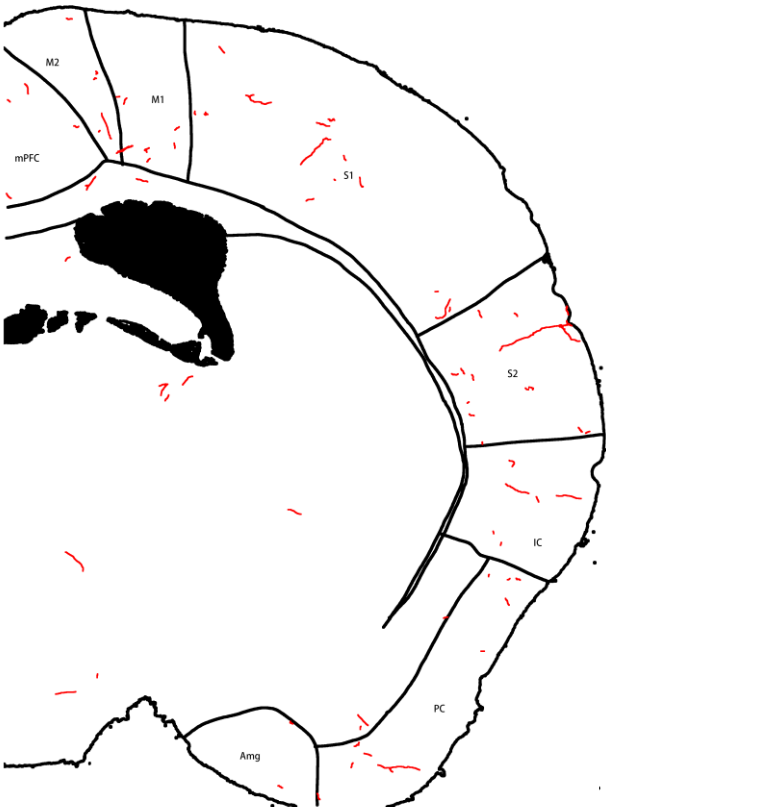

G: Bregma -1.39 mm

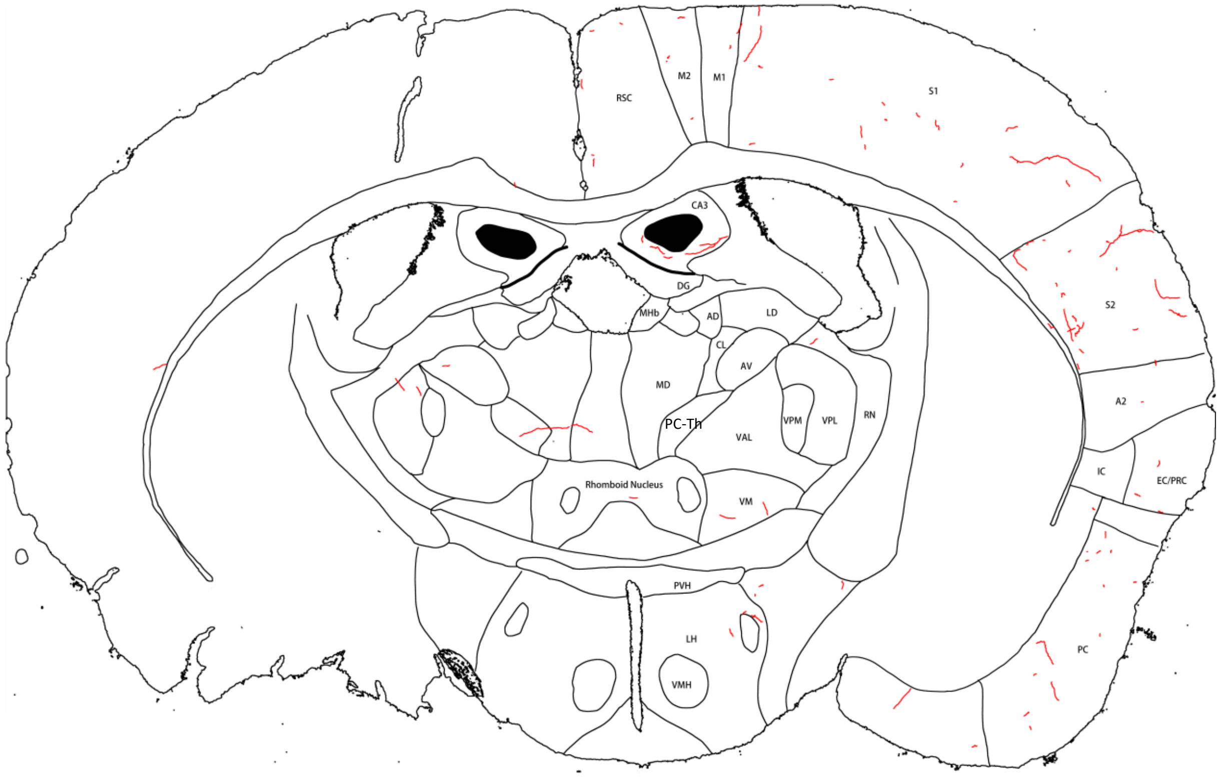


H: Bregma -1.49 mm

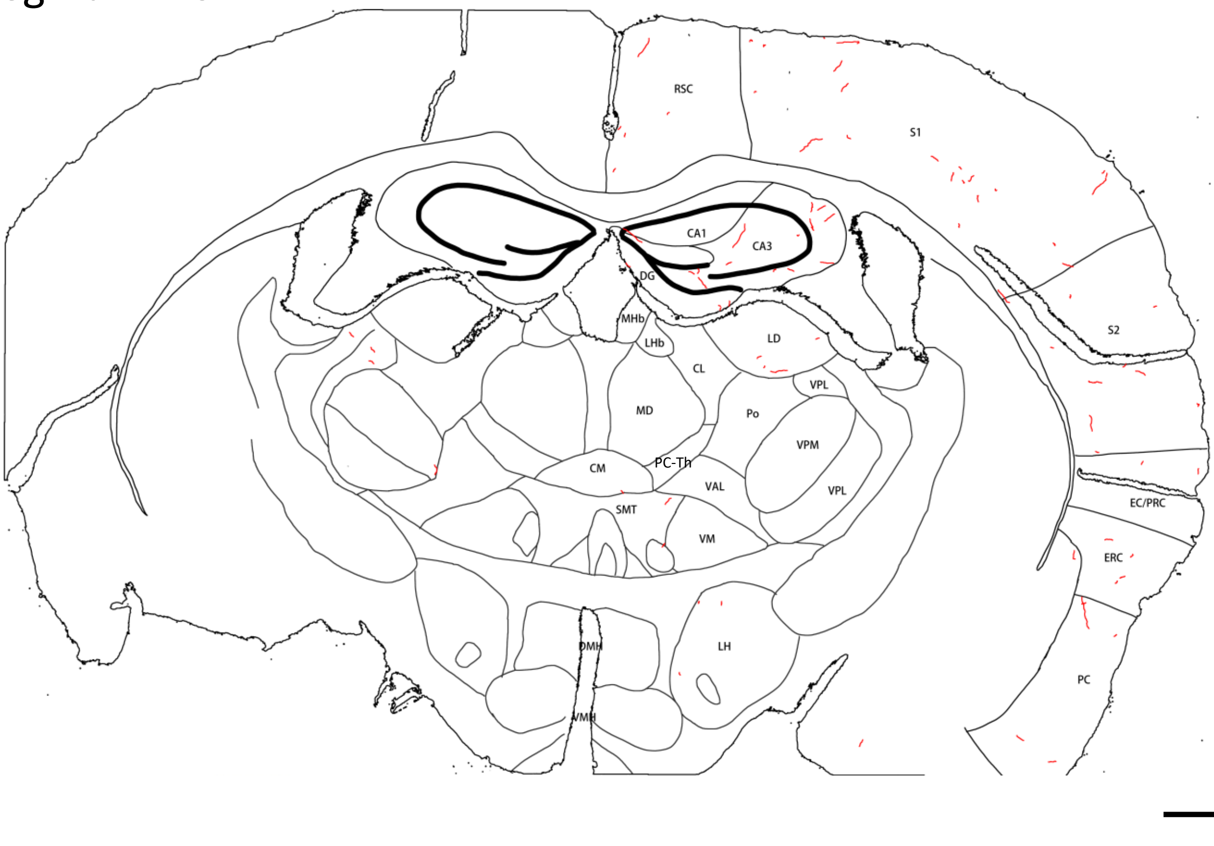

I: Bregma -1.82 mm

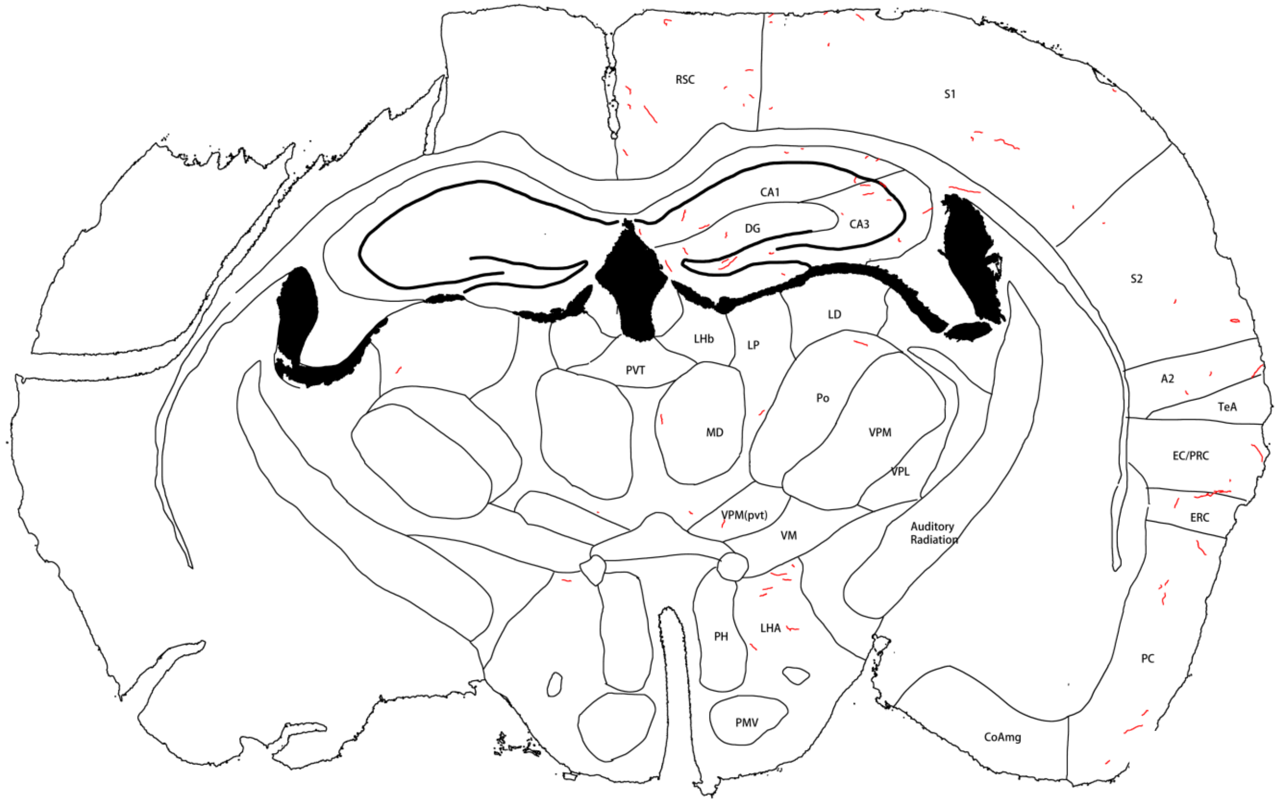


J: Bregma -2.03 mm

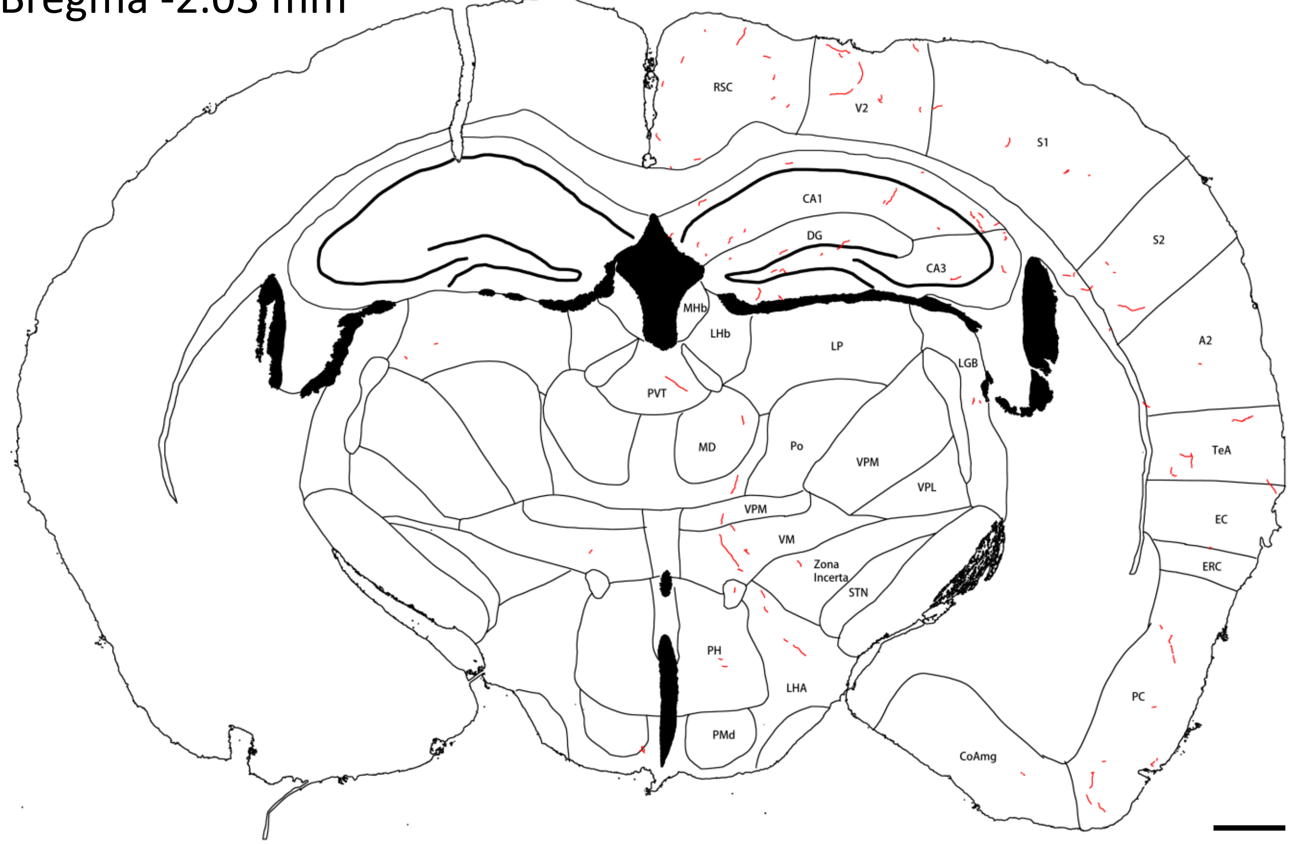

K: Bregma -2.93 mm

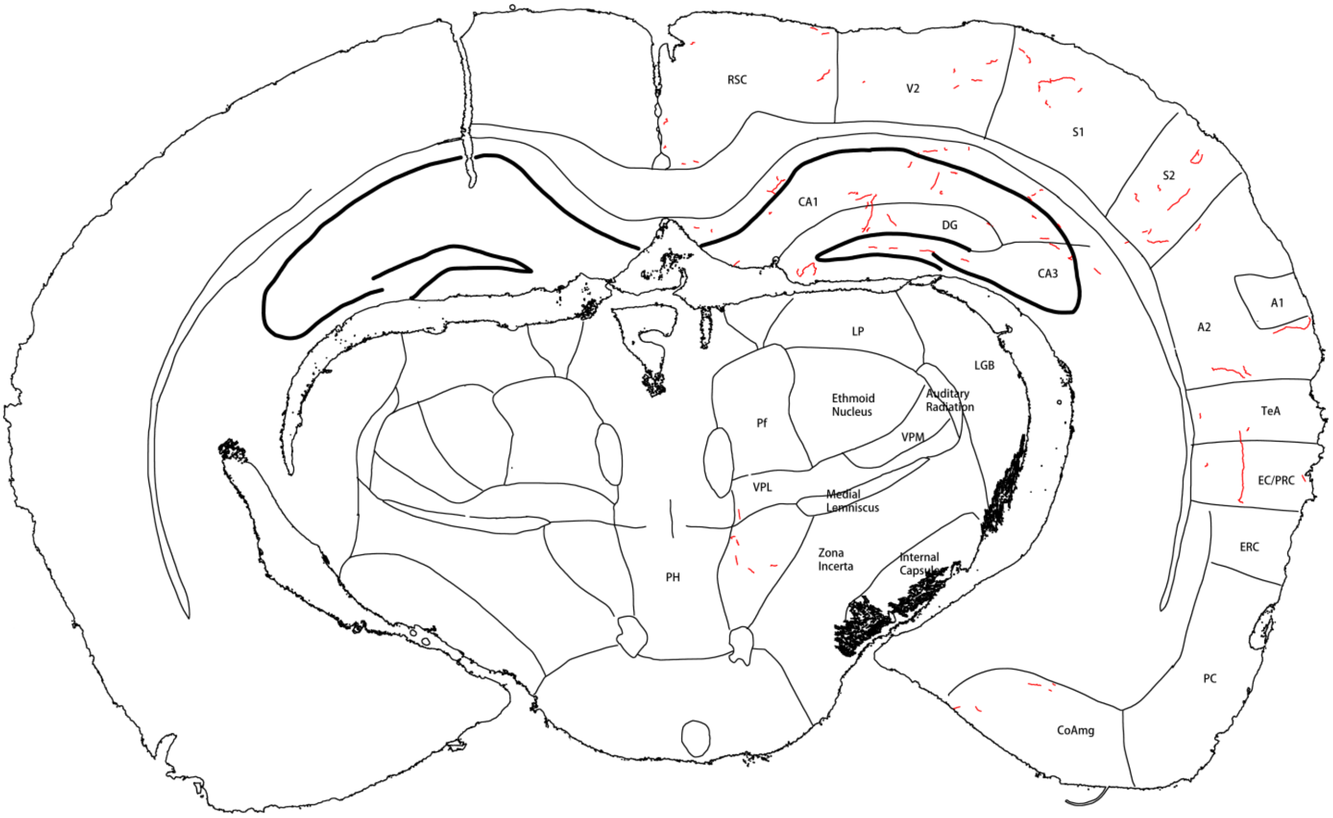


L: Bregma -3.17 mm

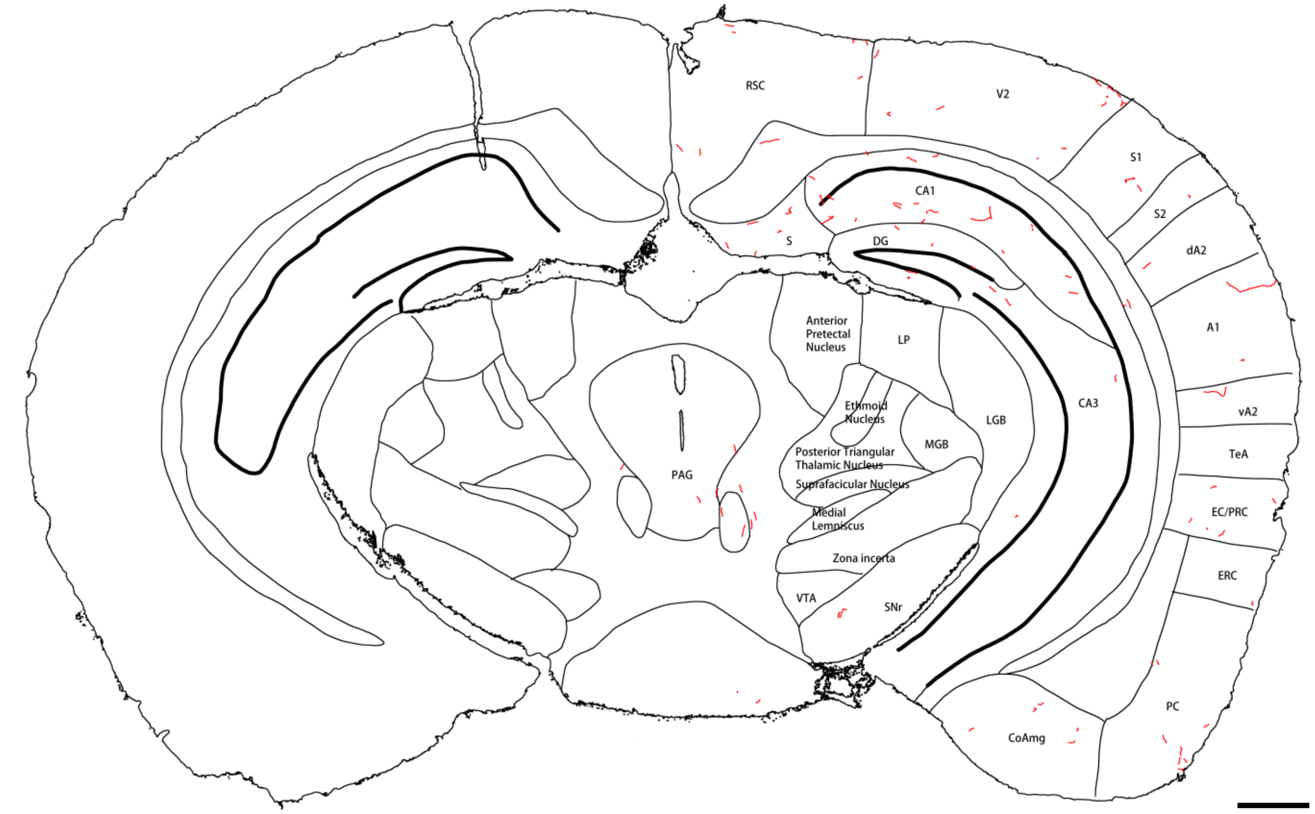

M: Bregma -2.26 mm

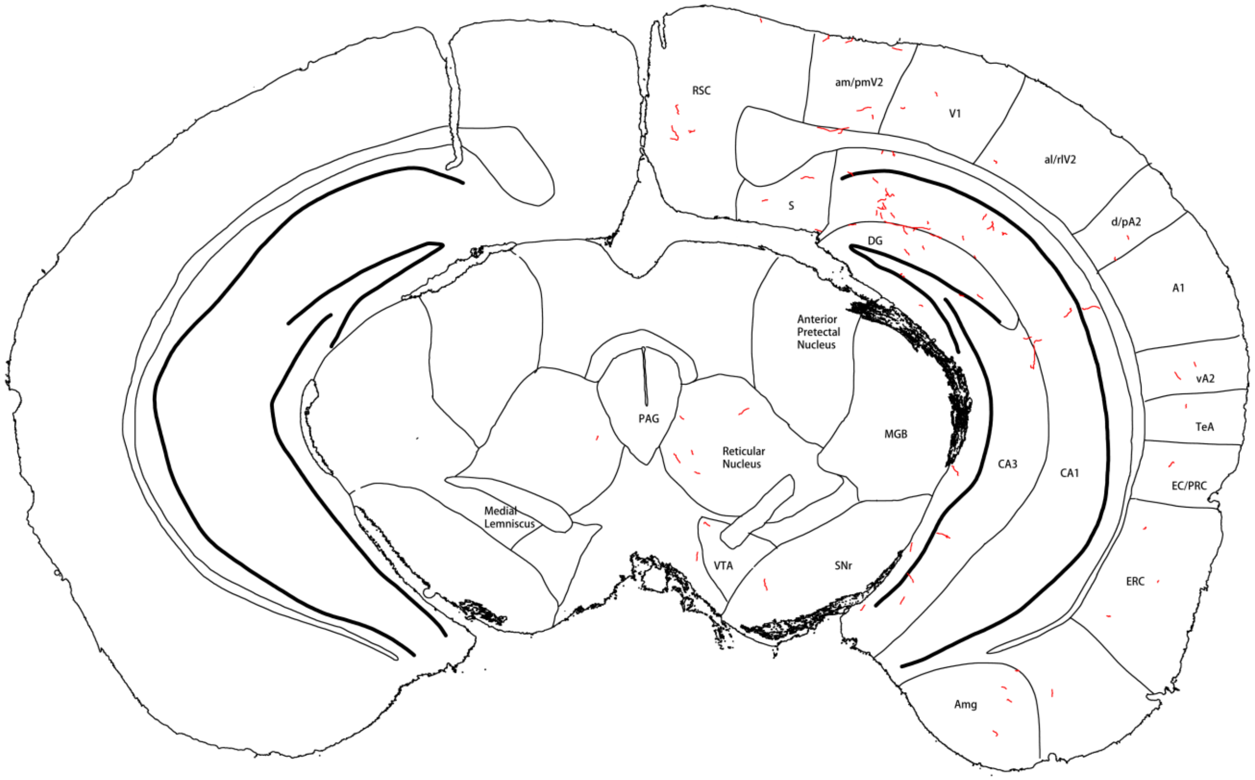


N: Bregma -2.62 mm

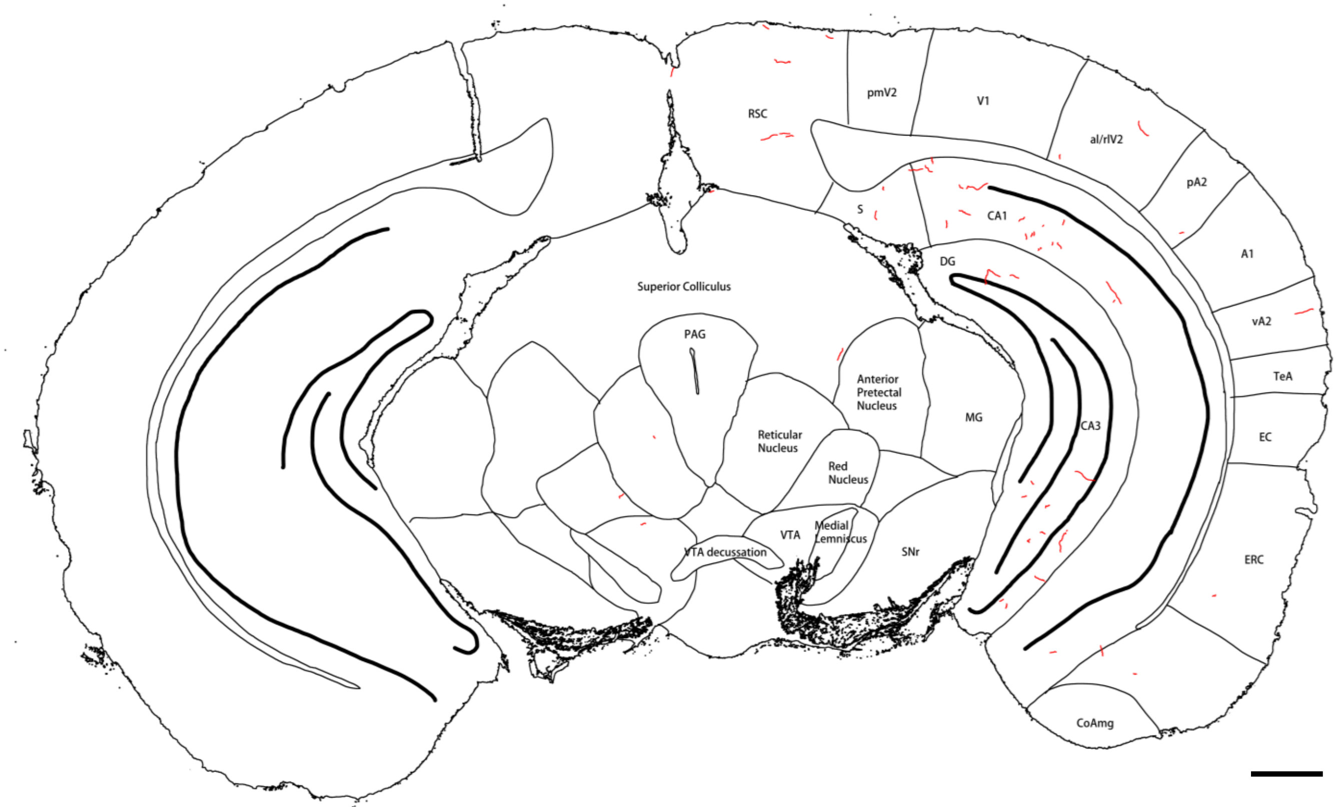

O: Bregma -3.40 mm

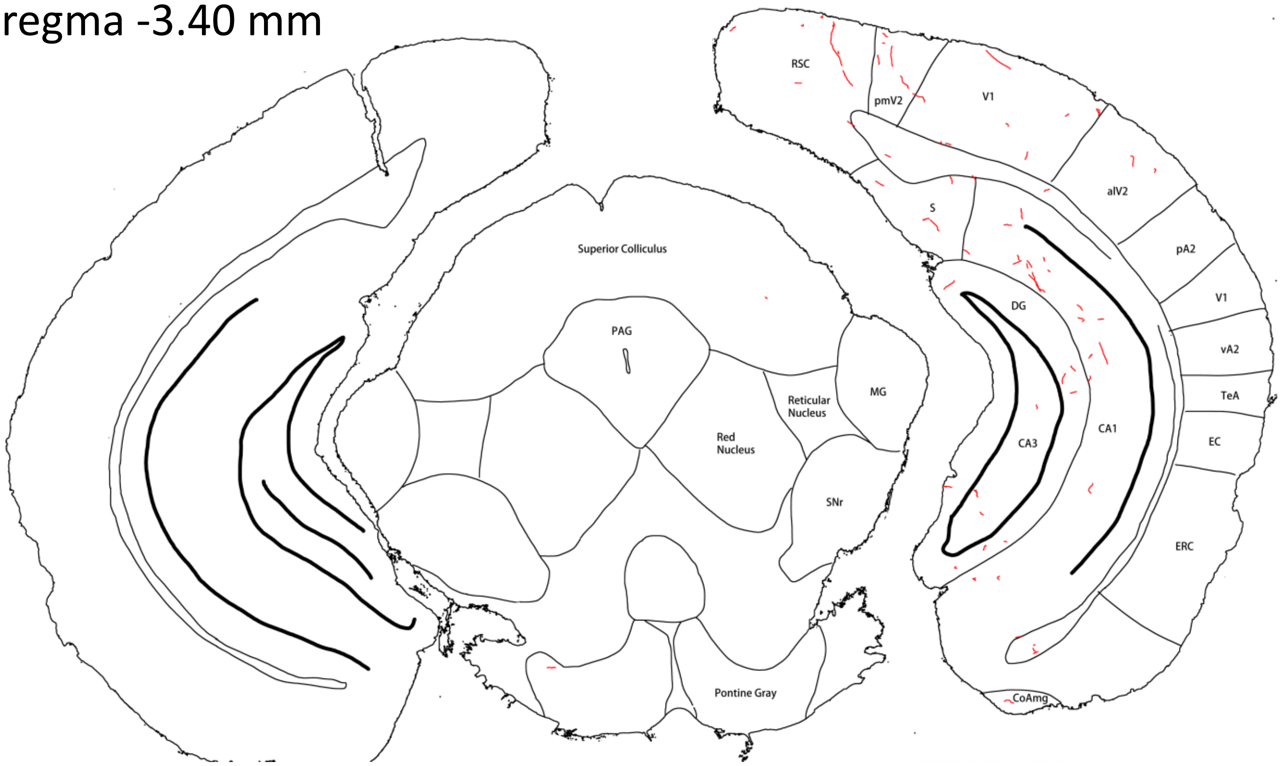


P: Bregma -3.65 mm

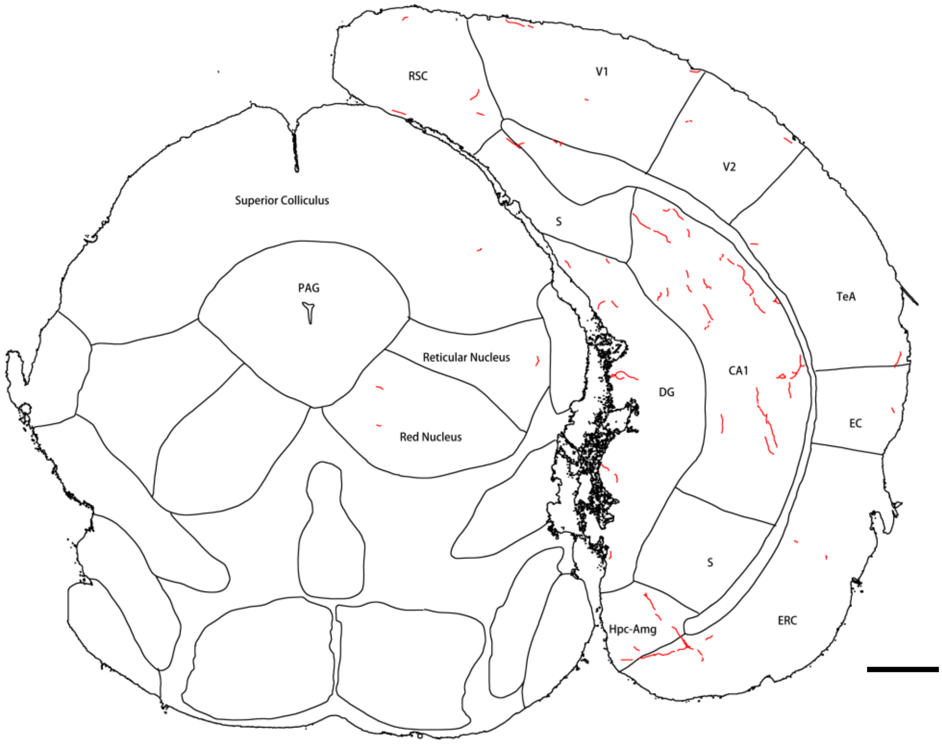


**Extended Figure 3C-3**. Tracing of NE fibers (red) in forebrain sections of **mouse #513**. Panels **A**-**F** are sections stained with FIHC; the others are sections stained with IHC/ABC. Yellow shadows in panels **A**-**D** mark the viral injection site with the core shown in panel **B**. Note that only sections where NE fibers are found are shown. Scale bar = 0.5 mm.
